# Recovering mixtures of fast diffusing states from short single particle trajectories

**DOI:** 10.1101/2021.05.03.442482

**Authors:** Alec Heckert, Liza Dahal, Robert Tjian, Xavier Darzacq

**Affiliations:** Department of Molecular and Cell Biology, Li Ka Shing Center for Biomedical and Health Sciences, University of California, Berkeley, CA 94720, USA; Eikon Therapeutics, Hayward, CA 94545, USA; Howard Hughes Medical Institute, Berkeley, CA 94720, USA; Department of Molecular and Cell Biology, Li Ka Shing Center for Biomedical and Health Sciences, CIRM Center of Excellence, University of California, Berkeley, CA 94720, USA

## Abstract

Single particle tracking (SPT) directly measures the dynamics of proteins in living cells and is a powerful tool to dissect molecular mechanisms of cellular regulation. Interpretation of SPT with fast-diffusing proteins in mammalian cells, however, is complicated by technical limitations imposed by fast image acquisition. These limitations include short trajectory length due to photobleaching and shallow depth of field, high localization error due to the low photon budget imposed by short integration times, and cell-to-cell variability. To address these issues, we developed methods to infer distributions of diffusion coefficients from SPT data with short trajectories, variable localization accuracy, and absence of prior knowledge about the number of underlying states. We discuss advantages and disadvantages of these approaches relative to other frameworks for SPT analysis.

**Significance statement:** Single particle tracking (SPT) uses fluorescent probes to track the motions of individual molecules inside living cells, providing biologists with a close view of the cell’s inner machinery at work. Commonly used SPT imaging approaches, however, result in fragmentation of trajectories into small pieces as the probes move through the microscope’s plane of focus. This makes it challenging to extract usable biological information. This paper describes a method to reconstruct an SPT target’s dynamic profile from these trajectory fragments. The method builds on previous approaches to provide information about challenging SPT targets without discrete dynamic states while accounting for some known biases, enabling observation of previously hidden features in mammalian SPT experiments.

## Introduction

Biological processes are driven by interactions between molecules. To understand the role of a molecular species in a process, a central challenge is to measure the subpopulations of the molecule engaged in distinct interactions without perturbing the living system’s steady state. Some interactions – such as complex formation or scaffold binding – are associated with changes in a molecule’s mobility. As a result, live cell single particle tracking (SPT), by separately observing the motion of individual molecules, is a promising tool to meet this challenge [1].

Advances in the past two decades have led to the application of live cell SPT beyond its original implementation on cellular membranes [2] [3] [4] [5]. These include the use of stochastic labeling to isolate a single emitter’s path [6], a principle that can be extended into intracellular settings with genetically encoded photoconvertible proteins [7] [8] or cell-permeable dyes [9] [10]. Another advance is stroboscopic illumination, which reduces blur associated with fast-diffusing emitters [11]. Together with modifications of TIRF microscopes [12], these innovations have enabled the application of SPT to intracellular settings with fast-moving subpopulations [13] [14] [15] [16]. We refer to this variant of SPT as stroboscopic photoactivated single particle tracking (spaSPT; Fig. 1A; Supplementary Movie 1).

**Figure 1:**
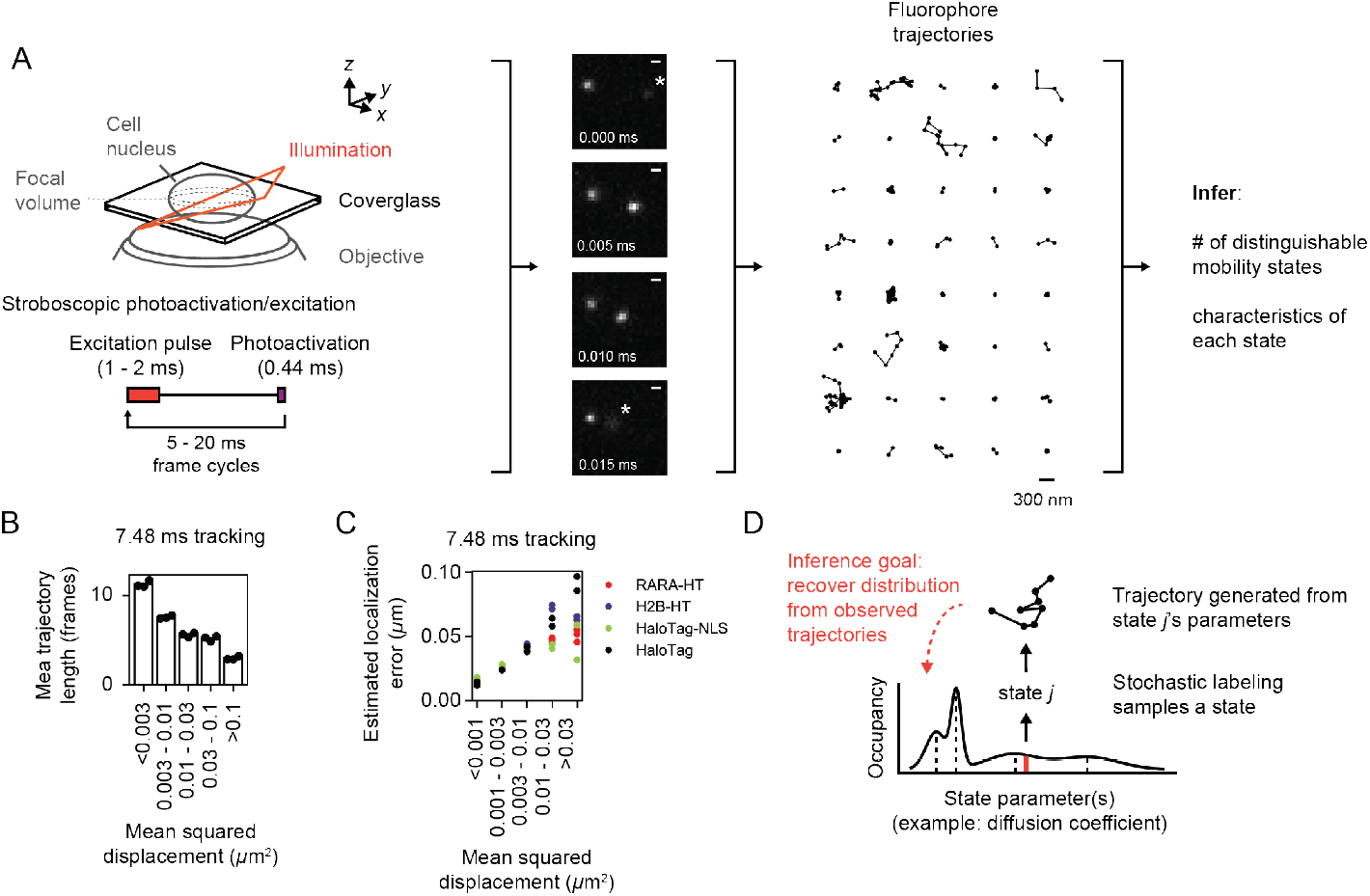
Fast stroboscopic single particle tracking (spaSPT). (A) Experimental setup; not to scale. An inclined illumination source is used in combination with a high-NA objective to resolve molecules in a thin slice in a cell. Laser photoactivation and excitation are pulsed to limit motion blur. After localization and tracking, the output is a set of short trajectories (mean track length 3-4 frames). Sample trajectories shown are from a 7.48 ms tracking movie with retinoic acid receptor *α*--HaloTag (RARA-HaloTag) labeled with photoactivatable JF549 in U2OS nuclei. The asterisks in the movie frames indicate particles that are either focalizing or defocalizing. (B) Influence of mobility on trajectory length. RARA-HaloTag trajectories from U2OS nuclei were classified into five categories based on their mean squared displacement (MSD). Individual data points represent the mean trajectory length of independent biological replicates using separate knock-in clones for RARA-HaloTag, and bar heights are the means across replicates. (C) Influence of mobility on localization error. Trajectories were categorized according to their MSD and for each category the localization error was estimated as the root negative covariance between subsequent jumps (see Supplementary Information). Individual data points represent biological replicates. (D) Schematic of the generative model for the inference frameworks in this manuscript. Each trajectory is assumed to represent a random draw from a distribution of state parameters, and the inference goal is to recover this distribution from the observed trajectories. Note that transitions between states are neglected in this framework.

Despite these advances, several problems remain when applying spaSPT to fast-moving emitters in 3D settings [17]. First, apparent motion in spaSPT originates both from the true motion of the emitter as well as localization error, the imprecision associated with our estimate for its position [18]. As in fixed cell super-resolution modalities [19] [20], localization error in spaSPT depends on the number of photons collected from an individual emitter per frame [21]. But spaSPT has an additional component of error due to *motion blur*, the convolution of the microscope’s 3D point spread function with the path of the emitter. Consequently, localization error in spaSPT depends on both the emitter’s mobility and its distance from the focus and is not trivial to measure [4] [22] [23]. Stroboscopic illumination reduces motion blur [11], but because camera integration times are never instantaneous, it cannot be removed entirely (Fig. 1C, S1B).

Second, the high numerical aperture objectives required to resolve single emitters induce short depths of field, typically less than a micron. Whereas bacteria such as *E. coli* are often small enough to fit into the resulting focal volume, mammalian cells – with depths 5-10 *µ*m or greater – cannot. As a result, intracellular SPT experiments only capture short transits of emitters through the focal volume with the duration of each transit related to the emitter’s mobility, a behavior termed defocalization (Fig. 1B, S1A, Supplementary Movies 2 and 3) [24] [25] [17]. This creates a sampling problem: slow particles with long residences inside the focal volume contribute a few long trajectories, while fast particles with short residences contribute many short trajectories. Mean trajectory length is often as little as 3-4 frames, severely limiting the ability to infer dynamic parameters (such as diffusion coefficient) from any single trajectory.

While methods for fast 3D imaging to mitigate this problem have been described [26] [27], they require higher photon budgets and are not yet applicable to fast-diffusing targets with high motion blur.

Third, the true number of dynamic subpopulations or “states” for a protein of interest is usually unknown *a priori*. Proteins often participate in many complexes with distinct dynamics. Model-dependent analyses that assume a fixed number of states [25] [16] [17], while powerful when combined with complementary measurements [14] [28], are limited to measuring coefficients of known models. To compound model complexity, a protein in one state may behave differently in distinct subcellular environments. Indeed, although spaSPT directly observes the spatial context for each trajectory [29], model-based analyses such as jump distribution modeling often discard this information.

The central problem for spaSPT analysis is to recover the underlying set of dynamic states for a protein target of interest, given a set of observed trajectories in the presence of these three obstacles.

To date, the most common approach to recover subpopulations from spaSPT data has been to construct histograms of the mean squared displacement (MSD), the maximum likelihood estimator for the diffusion coefficient in the absence of localization error. The MSD is highly variable for short trajectories and becomes especially error-prone when localization error is unknown [23]. More problematically, MSD histograms rely on the assumption that sampling from slow and fast states with equal fractional occupation produces the same number of trajectories, which leads to severe state biases in the presence of defocalization [25] [17]. Common preprocessing steps to select for long trajectories compound the problem by introducing biases for slow states that remain in focus.

A different approach to model selection is represented by vbSPT, a variational Bayesian framework for reaction-diffusion models that relies on the evidence lower bound to identify the number of states [30]. vbSPT excels at recovering occupations and transition rates for a small number of diffusing states, but it is not appropriate to apply in situations where the distribution of diffusion coefficients is not discrete and does not consider defocalization. As such, there is a need for methods that combine the flexibility of the MSD histogram approach with advantages of Bayesian methods like vbSPT, while accounting for biases induced by spaSPT imaging geometry.

Here, we examine two alternatives to MSD histograms useful for short trajectories. The first is based on a Dirichlet process mixture model (DPMM) and the second on a finite-state approximation to the DPMM that we refer to as a state array (SA). Exploring these techniques on simulated and real datasets, we find both DPMMs and SAs recover complex mixtures of states and can also be applied to non-discrete distributions of diffusion coefficients. While DPMMs outperform SAs when the localization error is known, SAs are more useful for experiments with limited prior knowledge about the error associated with each mobile state. Finally, we discuss limitations of these methods compared to other approaches.

## Results

### Two approaches to infer distributions of diffusion coefficients from spaSPT trajectories

As the target for inference, we considered a mixture of regular Brownian motions with localization error (RBMEs; SI Appendix A) enclosed in a spherical membrane with a thin focal volume bisecting the sphere, with dimensions resembling a mammalian cell nucleus (Fig. 2C). Each RBME is characterized by two parameters: a diffusion coefficient and a localization error magnitude. Emitters are subject to photoactivation and photobleaching throughout the sphere and are only observed when their positions coincide with the focal volume. Because no gaps are allowed during tracking, the result is a highly fragmented set of trajectories with mean length 3-5 frames. Simulation settings were chosen to approximate real spaSPT experiments, with bleaching rates ≥ 10 Hz, diffusion coefficients in the range 0-100 *µ*m^2^ s^*−*1^, and normally distributed localization error with standard deviation ≥ 30 nm. We sought to estimate the underlying distribution of diffusion coefficients from this data.

**Figure 2:**
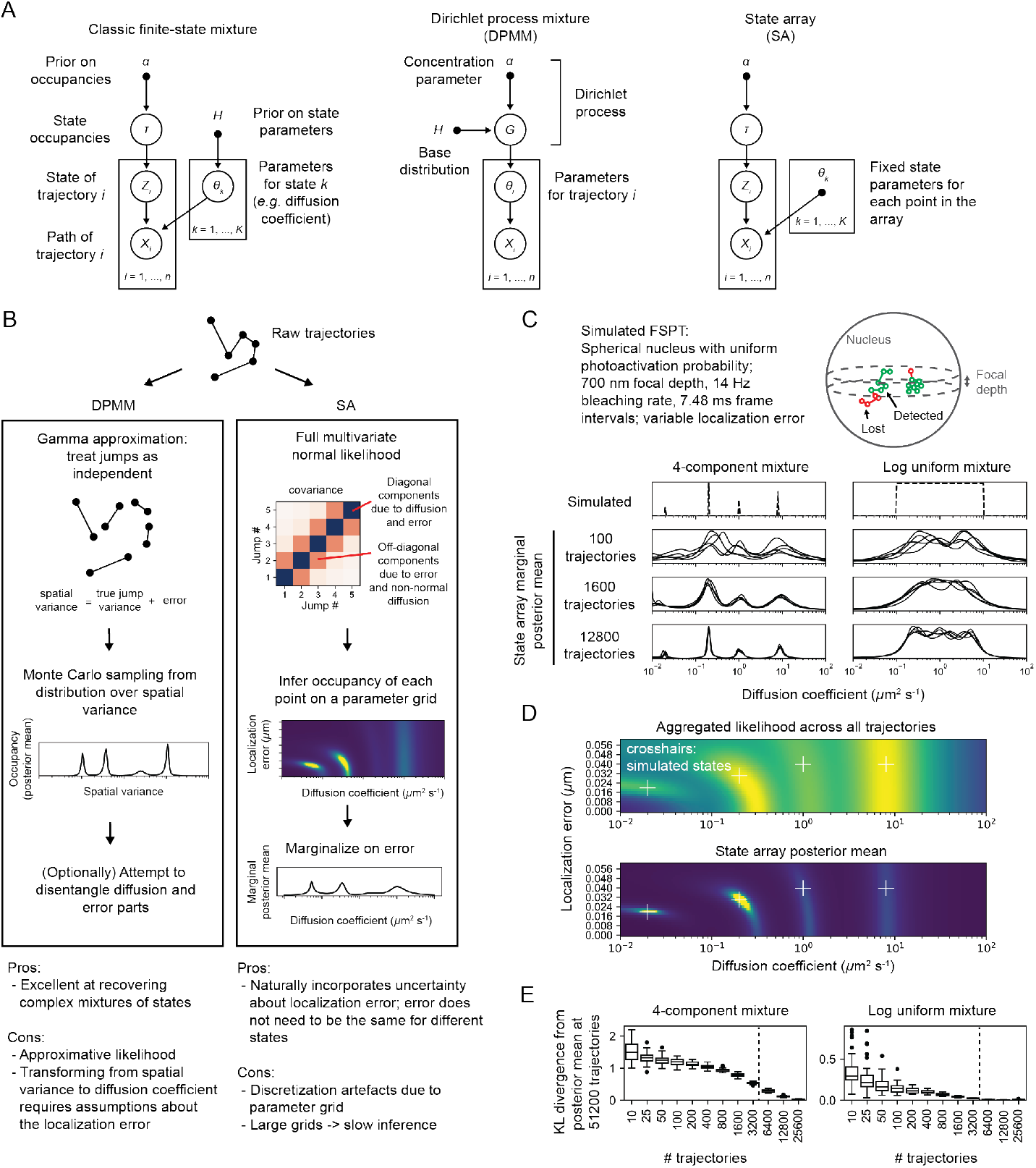
Schematic of Dirichlet process mixture models (DPMMs) and state arrays (SAs). (A) Graphical models for finite-state mixture models (FSMMs), DPMMs, and SAs. Open circles indicate random variables and solid circles indicate constants. (B) Schematic overview of the DPMM and SA methods applied to regular Brownian motion with localization error (RBME). (C) spaSPT simulations used to evaluate the performance of DPMMs and SAs. The dotted lines represent the simulated states, while the mean of the posterior distribution for DPMMs and SAs were used to estimate state occupations. (D) Comparison of the raw likelihood function against the posterior mean for mixtures of RBMEs. Crosshairs indicate the ground truth parameters for the four simulated states. (E) Convergence properties of the marginal posterior mean for SAs on the two kinds of simulations in (C). In these experiments, the posterior mean at 51200 trajectories was used as the distribution from which the KL divergence was calculated. The dotted line indicates the number of pseudocounts in the uniform prior on this SA grid.

An attractive approach to recover an arbitrary distribution of diffusion coefficients is the Dirichlet process mixture model (DPMM) [31]. DPMMs can be considered a modification of classic Bayesian finite-state mixture models (FSMMs) [32] (Supplementary Info). The two are compared as graphical models in Fig. 2A. Rather than inferring the occupations and parameters for each of *K* discrete states, DPMMs let *K*→ ∞ [33] and rely on the ability of Bayesian methods to identify sparse subsets of states sufficient to explain the observed trajectories. The priors and posteriors are continuous distributions over state parameters such as diffusion coefficient.

A challenge is that DPMM inference algorithms require evaluation of the model’s likelihood function for a large number of trajectory-state assignments at each iteration. When dealing with large datasets, costly likelihood functions, and non-conjugate priors – for instance, uniform priors over the diffusion coefficient and localization error – this becomes computationally expensive [34].

The approaches in this manuscript represent two responses to this problem. First, we considered a DPMM with an approximative likelihood function that is much faster to compute, obtained by treating the RBME as a Markov process (Fig. 2B, left) [35]. This assumption is strictly true only when localization error is zero and is the same assumption that accompanies the use of the MSD as the maximum likelihood estimator for the diffusion coefficient [23]. To estimate the posterior distribution, we use a Markov chain Monte Carlo approach related to Neal’s Algorithm 8 [36].

The second approach is to replace the DPMM with a conceptually similar method we call a state array (SA). An SA is a grid of state parameters that span some target part of parameter space – for instance, a range of diffusion coefficients. Each point in this grid is treated as a separate state. Because the parameters for each state are fixed, likelihoods for each trajectory-state assignment only need to be evaluated once; the sole inferential goal is then to calculate the posterior occupations of each point in the grid. As a result, SAs can handle more complex likelihood functions that incorporate localization error. By marginalizing the posterior distribution on localization error, the method naturally incorporates uncertainty about the localization error of different states.

The primary drawbacks of SAs are the discretization artefacts expected to result when the spacing of the parameter grid is chosen too coarsely with respect to local changes in the likelihood. (DPMMs are obtained by letting the grid spacing go to zero.) SAs have conceptual similarities to an approach recently proposed to measure the distribution of chromatin binding residence times for transcription factor SPT [37], with a variational Bayesian inference routine [38] substituting for the inverse Laplace transform. Both DPMMs and SAs are described in detail in the Supplementary Information.

### Evaluating DPMMs and SAs on simulated spaSPT data

First, we compared the efficacy of DPMMs, SAs, and the MSD histogram approach on simulated mixtures of diffusing states (Fig. 3, S2). We divided these simulations into three classes with increasing difficulty. In the first class, localization error for all states was provided as a known constant to the algorithms (Fig. S2A). In the second class, localization error was held constant for all states but was unknown to the algorithms (Fig. S2B). In the third class, localization error was allowed to vary between diffusive states and was also unknown to the algorithms (Fig. S2C).

**Figure 3:**
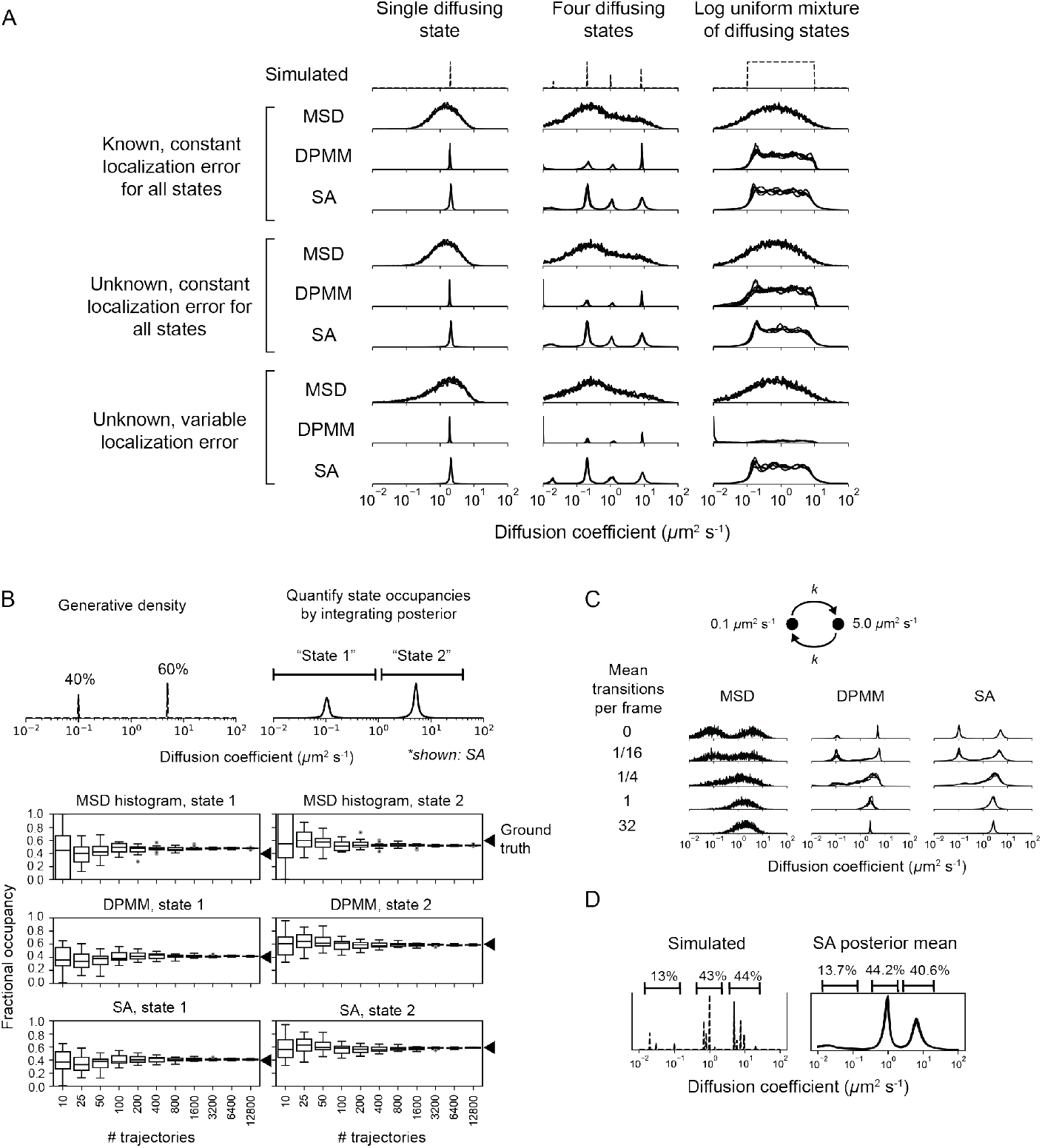
Comparison of DPMMs, SAs, and MSD histograms on simulated spaSPT. (A) Different combinations of diffusing states were simulated in a thin (700 nm) focal volume with 7.48 ms frame intervals. Simulations were divided into three classes: (i) constant (30 nm) localization error which was provided to the inference algorithms, (ii) constant (30 nm) localization error which was unknown to the algorithms, and (iii) localization error that varied with the diffusion coefficient and was unknown to the algorithms. For DPMM and SA, the mean of the posterior distribution is shown. (B) Fractional occupation was estimated by integrating the normalized histogram (for the MSD method) or by integrating the mean of the posterior distribution (for DPMM and SA methods). The limits of integration for each state were determined based on visual inspection of the posterior distribution, and were set to 0 - 1 *µ*m^2^ s^*−*1^ for the first state and 1 - 40 *µ*m^2^ s^*−*1^ for the second state. (C) Effect of state transitions on the MSD, DPMM, and SA approaches. Two diffusing states with first-order transitions were simulated, and the transition rate constant was varied. (D) Inferring mixtures of diffusing states with similar diffusion coefficients. Occupations as percentages were obtained by integrating the indicated parts of the distribution.

DPMMs and SAs both recovered the distribution of diffusion coefficients for simulations in class 1 with a resolution that exceeded the MSD histogram approach (Fig. 3, S2A). With large samples of trajectories, DPMMs and SAs inferred even non-discrete distributions of states. Notably, in the presence of multiple diffusing states with similar diffusion coefficients, both DPMMs and SAs tended to identify a single state with occupation equal to the sum of the occupations for each true state (Fig. 3D, S5B).

The DPMM approach was more precise than the SA approach on simulations in class 1. In contrast, when knowledge of the localization error was removed, the SA approach outperformed both the MSD and DPMM approaches. Indeed, assuming a constant localization error had strong effects on the DPMM’s ability to infer diffusion coefficients for slow-diffusing states with variable localization error, while SAs were essentially unperturbed by variations in the localization error (Fig. S3). Because localization error is frequently difficult to measure in spaSPT, this quality of SAs is useful.

We also compared the accuracy of state occupation estimates obtained from the three approaches (Fig. 3B, S4). As the number of trajectories used in inference increased, the occupations estimated by DPMMs and SAs converged to values within 1% of the true underlying occupations. In contrast, the MSD approach was associated with large systematic errors due to the fact that the approach counts by trajectories rather than jumps and does not account for defocalization, an effect previously reported [25] [17]. Importantly, when states with diffusion coefficients outside the support were used, DPMMs and SAs still accurately recovered state occupations by using the closest diffusion coefficient available in the support (Fig. S5A).

A central limitation of DPMMs and SAs is that they do not account for transitions between diffusive states. To determine the effect of state transitions on the output of these algorithms, we simulated mixtures of two diffusive states with increasing transition rates (Fig. 3C, S6). While slow transition rates have a negligible effect on the estimated state profile, transition rates approaching the frame interval result in the inference of a single apparent state with intermediate diffusion coefficient (Fig. S6C), consonant with a well-known result from reaction-diffusion systems [39]. The shift from the two-state to single-state regime occurs in a narrow window of mean state dwell times between 0.05 and 0.5 frame intervals. Consequently, performing multiple spaSPT experiments with different frame intervals may be a useful way to identify fast state transitions for proteins with complex dynamic profiles.

### Performance of DPMMs and SAs on experimental spaSPT

To examine the performance of SAs on real data, we acquired a spaSPT dataset in U2OS osteosarcoma nuclei with endogenously tagged retinoic acid receptor--HaloTag (RARA-HT) (Fig. S7) [40] [41]. RARA-HT is a type II nuclear receptor that heterodimerizes via its ligand-binding domain (LBD) with the retinoid X receptor (RXR) to form a complex competent to bind chromatin and regulate target genes [42] [43] [44] [45] [46] [47] [48] (reviewed in [49]). In addition, association of coregulator complexes with the RAR/RXR heterodimer has been shown to influence the dimer’s dynamics in FCS studies [50] [51]. As such, RARA-HT is expected to inhabit a variety of dynamic states in spaSPT.

For comparison, we also performed identical spaSPT experiments with histone H2B-HaloTag (H2B-HT), a protein with a high-occupation immobile state [16] [52], as well as HaloTag and HaloTag-NLS (HT and HT-NLS), which are fast-diffusing proteins with low immobile fractions.

The four proteins presented distinct dynamic profiles (Fig. 4A). For both HT and HT-NLS, the SA identified a single highly mobile state. In agreement with previous reports [29], we observed that addition of the NLS reduces HaloTag’s diffusion coefficient by two- to three-fold. In contrast, both RARA-HT and H2B-HT had substantial immobile fractions, accounting for roughly 40% and 70% of their total populations respectively (Fig. 4C). The mobile subpopulations for RARA-HT and H2B-HT differed markedly. Whereas H2B-HT presented a fast population at 8-10 *µ*m^2^ s^−1^, RARA-HT inhabited a broad spectrum of diffusing states ranging from 0.3 to 10.0 *µ*m^2^ s^−1^. Biological replicates gave similar results (Fig. S8A).

**Figure 4:**
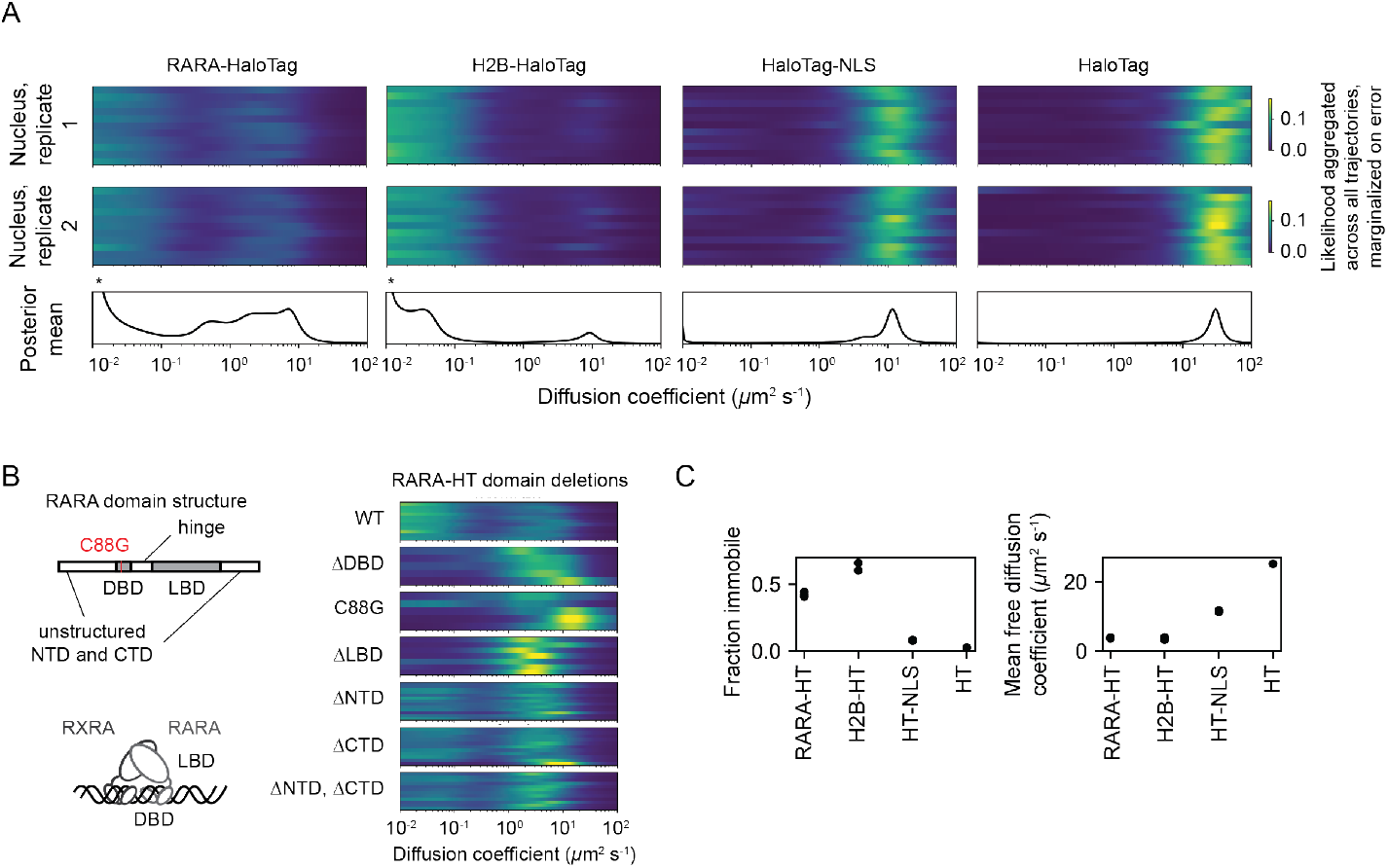
Using state arrays to recover subpopulations from experimental spaSPT. All spaSPT experiments were performed with the photactivatable dye PA-JFX549 using a TIRF microscope with HiLo illumination, 7.48 ms frame intervals, and 1 ms excitation pulses. (A) Posterior mean occupations for four different tracking targets, compared to the raw likelihood function. The upper two panels are the RBME likelihood function aggregated across all trajectories, weighted by the number of jumps in each trajectory and marginalized on the localization error component. The bottom panel is the posterior for a run of the SA algorithm. Asterisks for RARA-HaloTag and H2B-HaloTag indicate that the immobile fraction for these constructs has been truncated to visualize the faster-moving states. (B) Aggregate likelihood functions for RARA-HaloTag constructs bearing domain deletions or point mutations. (C) Quantification of the immobile fractions and mean free diffusion coefficients for the four constructs in (A). The “immobile fraction” was defined as the total occupation below 0.05 *µ*m^2^ s^*−*1^, while the mean free diffusion coefficient was the occupation-weighted mean of the diffusion coefficients above this threshold.

To determine the origins of the dynamic states observed for RARA-HT, we performed domain deletions (Fig. 4B). Removal of either the DNA-binding domain (DBD) or LBD resulted in loss of the immobile population. Because both the DBD and LBD are required for chromatin binding by the RAR/RXR heterodimer, this suggests that the immobile fraction represents chromatin-bound molecules. To confirm this, we introduced a point mutation (C88G) in the zinc fingers for the RARA-HT DBD that abolishes DNA-binding in vitro [53]. This led to loss of the immobile fraction (Fig. 4B). Deletion of the unstructured N-terminal domain (NTD) or C-terminal domain (CTD) had a milder effect, suggesting that these domains are not the primary determinants of the dynamic behavior of RARA-HT.

To understand the origins of heterogeneity in the diffusive profile, we performed three variants of bootstrap aggregation (Fig. S8B). The primary origins of variability for both DPMMs and SAs were cell-to-cell rather than clone-to-clone variability or intrinsic variability due to finite sample sizes.

### Spatiotemporal context of cellular protein dynamics

The full posterior model for DPMMs and SAs involves probability distributions over the diffusion coefficient and/or localization error for every trajectory in a spaSPT dataset. Because spaSPT trajectories are short, a single trajectory cannot provide high-confidence information about state parameters. But aggregating information across trajectories in the posterior distribution offers a potential route to understand how the diffusion coefficient varies spatially and temporally within a dataset.

We explored this aspect of the SA method on a U2OS nucleophosmin-HaloTag (NPM1-HT) spaSPT dataset. NPM1-HT exhibits partial nucleolar localization (Fig. S9B) and distinct dynamic behavior inside and outside nucleoli [54]. The SA method identified a broad range of diffusion coefficients for NPM1-HT, with three modes including an effectively immobile population (Fig. 5A). Selecting four regions of this spectrum for analysis, we visualized the posterior likelihood as a function of space (Fig. 5B). Evaluating the local posterior mean also provided a way to measure local occupations for each of these states (Fig. 5B, S9C). This analysis revealed that some populations (including a slow-moving mobile state at 0.23 µm2 s-1) are enriched in nucleoli, while others (for instance, a fast-moving state at 4 µm2 s-1) are depleted and still others show no preference (Fig. 5C). Notably, these preferences are apparent even in the raw likelihood function for trajectories in each compartment (Fig. S9D).

**Figure 5:**
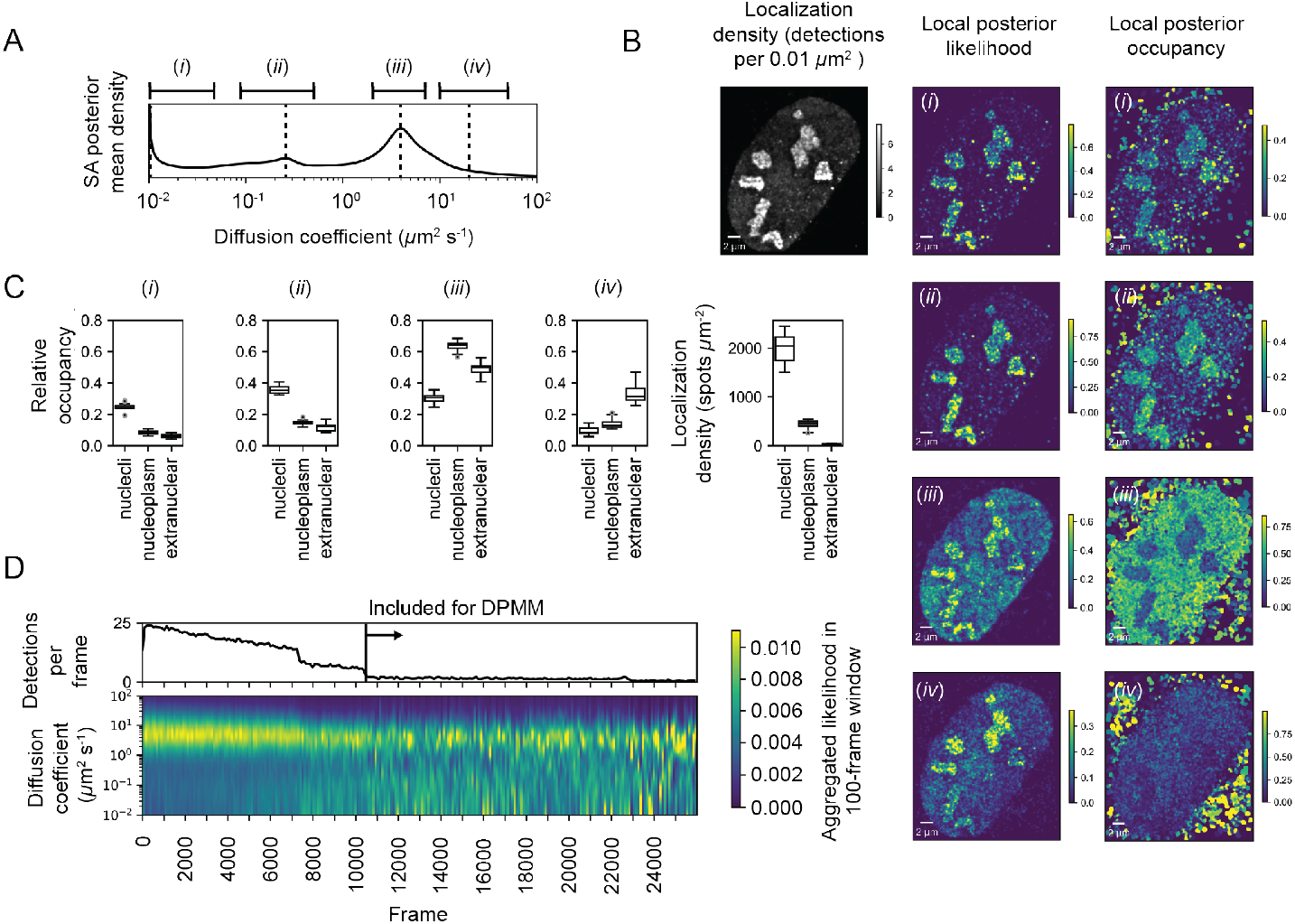
Applications of spatially and temporally indexed posterior state occupations. (A) Posterior mean occupations for a state array evaluated on NPM1-HaloTag trajectories in U2OS nuclei. The dotted lines indicate discrete diffusion coefficients that were isolated for analysis in subsequent panels. (B) Sample spatially-resolved likelihoods under the posterior model in (A) for NPM1-HaloTag trajectories in a U2OS nucleus. The posterior model over the diffusion coefficient was evaluated for each of the origin trajectories, and these points were then used to perform a kernel density estimate with a 100 nm Gaussian kernel. For the local normalized occupation, these KDEs were normalized to estimate the relative fractions of molecules in each state. (C) Quantification of the results in (B) for 15 nuclei. “Nucleoplasmic” trajectories were defined as trajectories outside nucleoli but inside the nucleus. (D) Example usage of the likelihood function to assess the effect of localization density on apparent state occupations. The RBME likelihood was aggregated across trajectories in 100 frame windows, then marginalized on localization error. Note the slight decrease in the apparent diffusion coefficient of population (iii) and the increase in the immobile population upon the decrease in localization density, probably reflecting tracking errors.

The NPM1-HT tracking experiments were performed with an acquisition sequence comprising several phases with distinct levels of photoactivation. As a result, the localization density varied temporally in each movie. To understand the effect of localization density on the diffusion coefficient likelihoods, we aggregated trajectory likelihoods in 100 frame temporal blocks (Fig. 5D). These experiments demonstrated that high localization densities led to a deflation in the occupation of slower-moving states, probably due to tracking errors. As a result, only phases with low localization density were used for posterior estimation. This demonstrates how the temporal indexing of the posterior likelihood function for trajectories can be used to guide subsequent analysis.

## Discussion

spaSPT with fast-diffusing proteins in 3D presents unique challenges for data analysis. In particular, the issues of state bias arising from imaging geometry, limited information available from any single trajectory, and variable localization error must be addressed prior to biological interpretation of spaSPT data.

Methods based on least-squares fitting of the jump length CDF have interpreted spaSPT data with two- and three-state models [25] [17] but extend poorly to more complex models due to overfitting and do not provide a way to distinguish between competing models. In contrast, nonparametric methods like the MSD histogram could in theory be used to identify any dynamic profile for any number of states, but in practice have poor resolution due to short trajectory length and also introduce state biases by assuming that slow and fast states with equal occupation present the same number of trajectories [17].

The two methods discussed here, DPMMs and SAs, occupy a middle ground and accurately identify state occupations in spaSPT data while avoiding assumptions about the number of states. These methods borrow from the capability of Bayesian methods to identify sparse explanatory models from more complex alternatives.

As an alternative to SAs, a DPMM for the full RBME likelihood (evaluating a posterior over the continuous state space of both diffusion coefficient and localization error) could be used to identify mixtures of RBMEs. While this approach is feasible, it is slow due to the requirement for evaluation of the RBME likelihood for a large number of trajectory-state assignments at each iteration of the algorithm. The gamma likelihood based DPMM and SAs considered here are approximations to the full DPMM that are accurate and fast enough for practical use without resorting to specialized implementations. SAs, by naturally incorporating uncertainty about localization error, are particularly useful for spaSPT datasets in which the localization error can vary between individual movies.

DPMMs and SAs have several limitations. DPMMs require prior measurement of the localization error, while SAs require selection of a parameter grid with spacing fine enough to avoid discretization artefacts. In addition, some limitations are shared by both DPMMs and SAs. These include the fact that neither method considers state transitions, and both share the inability to distinguish localization error from non-Brownian diffusion (e.g. subdiffusion) because both of these manifest primarily in the off-diagonal components of the RBME covariance matrix.

Neither DPMMs nor SAs have any kind of built-in mechanism to distinguish true jumps from tracking errors; they both rely on trajectories produced by another algorithm. It may be possible to combine both tracking and state occupation estimation into a single Bayesian inference routine in which the posterior over diffusion coefficients, localization error, and occupations for each state is jointly evaluated with a posterior over possible connections between particles.

## Acknowledgments

We would like to thank Luke Lavis for generously synthesizing the Janelia Fluor dyes used in these experiments, to Thomas Graham for his creative suggestions about the defocalization problem that led to the algorithm introduced in SI Appendix B, to Ana Robles for the monumental task of keeping our SPT microscopes in working order, and to Anders Hansen and Maxime Woringer for insights and advice at the outset of the project. Claudia Cattoglio provided indispensble advice on Western blots. Portions of this work were performed on shared instrumentation at the UC Berkeley Cancer Research Laboratory Molecular Imaging Center (MIC), supported by The Gordon and Betty Moore Foundation. We thank MIC gurus Holly Aaron and Feather Ives for their assistance with shared microscopes. Sanger sequencing was performed at the UC Berkeley DNA Sequencing Facility. This work was supported by NIH grant 1U54CA231641 (X. D.) and by the Howard Hughes Medical Institute (CC34430, R. T.). A. H. was supported by the NIH Stem Cell Biological Engineering pre-doctoral fellowship T32 GM098218. We would like to thank David Schaffer for his support and feedback throughout the project as a program director for the T32 fellowship.

## Materials and methods

### Plasmids

Unless otherwise noted, all PCRs were performed with New England Biosciences Phusion High-Fidelity DNA polymerase (M0530S) and Gibson assemblies [55] were performed with New England Biosciences Gibson Assembly Master Mix (E2611S) following manufacturer’s instructions. Cloning and expression of plasmids was performed in *E. coli* DH5*α* using the Inoue protocol [56]. Plasmids used for nucleofections were purified by Zymo midiprep kit (Zymo D4200) and concentrations were quantified by absorption at 260 nm. Cloning primers were synthesized by Integrated DNA Technologies as 25 nmol DNA oligos with standard desalting, and sequences were verified by Sanger sequencing at the UC Berkeley DNA Sequencing Facility.

We produced the vector PB PGKp-PuroR L30p MCS-GDGAGLIN-HaloTag-3xFLAG by amplifying the human L30 promoter with prAH675 and prAH676 and assembling into AsiSI- (NEB R0630) and XbaI- (NEB R0145) digested PB PGKp-PuroR EF1a MCS-GDGAGLIN-HaloTag-3xFLAG. For the expression plasmid PB PGKp-PuroR EF1a 3x-FLAG-HaloTag-GDGAGLIN, we cloned three tandem copies of the SV40 nuclear localization sequence into XbaI- and BamHI-HF (NEB R3136)-digested PB PGKp-PuroR EF1a 3xFLAG-HaloTag-MCS using Gibson assembly.

For constructs expressing RARA-HaloTag domain deletions and point mutations, we first cloned the RARA coding sequence out of U2OS cDNA by extracting RNA from cycling U2OS cells with a Qiagen RNeasy kit (Qiagen 74104), preparing cDNA with the iScript Reverse Transcription Supermix (Bio-Rad 1708840), amplifying the CDS with prAH495 and prAH496, then assembling into an XbaI- and NotI-HF- (NEB R3189) digested PB PGKp-PuroR EF1a MCS-GDGAGLIN-HaloTag-3xFLAG using Gibson assembly. Next, to produce the mutants, we amplified parts of the RARA coding sequence in PCR fragments while introducing point mutations or domain deletions at the intersections of the fragments. PCR fragments were assembled into XbaI- and BamHI-HF-digested PB PGKp-PuroR L30p-MCS-GDGAGLIN-HaloTag-3xFLAG using Gibson assembly. The primers used for each construct were as follows: for PB PGKp-PuroR EF1a RARA[NTD]-HaloTag-GDGAGLIN-3xFLAG, PCR fragment 1 was produced with prAH1111 and prAH1112; for PB PGKp-PuroR EF1a RARA[CTD]-HaloTag-GDGAGLIN-3xFLAG, PCR fragment 1 was produced with prAH1113 and prAH1114; for PB PGKp-PuroR EF1a RARA[NTD, CTD]-HaloTag-GDGAGLIN-3xFLAG, PCR fragment 1 was produced with prAH1111 and prAH1114; for PB PGKp-PuroR EF1a RARA[C88G]-HaloTag-GDGAGLIN-3xFLAG, PCR fragment 1 was produced with prAH1113 and prAH1069 and PCR fragment 2 was produced with prAH1112 and prAH1070; for PB PGKp-PuroR EF1a RARA[DBD]-HaloTag-GDGAGLIN-3xFLAG, PCR fragment 1 was produced with prAH596 and prAH704 and PCR fragment 2 was produced with prAH597 and prAH705; for PB PGKp-PuroR EF1a RARA[LBD]-HaloTag-GDGAGLIN-3xFLAG, PCR fragment 1 was produced with prAH596 and prAH706 and PCR fragment 2 was produced with prAH597 and prAH707.

To generate the plasmid-based homology repair donor for gene editing at the human RARA exon 9 locus, we assembled the following fragments by Gibson assembly. For fragment 1, we digested the pUC57 vector with EcoRI and HindIII. For fragment 2, we amplified the left homology arm out of U2OS genomic DNA with prAH599 and prAH600. For fragment 3, we amplified the GDGAGLIN-HaloTag-3xFLAG insert out of the plasmid PB PGKp-PuroR L30p MCS-GDGAGLIN-HaloTag-3xFLAG with prAH601 and prAH602. For fragment 4, we amplified the right homology arm out of U2OS genomic DNA with prAH603 and prAH604.

To generate guide RNA/Cas9 expression plasmids for gene editing at the human RARA exon 9 locus, we cloned the two guide RNA sequences under a U6 promoter in a vector that coexpresses the sgRNA, mVenus, and *S. pyogenes* Cas9, which has been previously described [16].

For luciferase assays, we used the retinoic acid-responsive firefly luciferase expression vector pGL3-RARE-luciferase (Addgene plasmid #13458; http://n2t.net/ad-dgene:13458; RRID:Addgene_13458), a gift from T. Michael Underhill [57]. Renilla luciferase was expressed from pRL CMV Renilla (Promega E2261).

### Tissue culture

Human U2OS cells (female, 15 year old, osteosarcoma) obtained from the UC Berkeley Cell Culture Facility were cultured under 5% CO_2_ at 37 degrees C in DMEM containing 4.5 g/L glucose supplemented with 10% fetal bovine serum and 10 U/mL penicillin-streptomycin. Cells were subpassaged at a ratio of 1:6 every 3-4 days. The stable cell line expressing H2B-HaloTag-SNAPf was described previously [16] [52]. Expression of HaloTag, HaloTag-NLS, and point mutants and domain deletions of RARA-HaloTag were induced by nucleofection of PiggyBac vectors containing the proteins under EF1a promoters. Expression of wildtype RARA-HaloTag and NPM1-HaloTag were induced by endogenous gene editing, as described in the “CRISPR/Cas9-mediated gene editing” section.

For spaSPT experiments, cells were grown on 25 mm circular No. 1.5H cover-glasses (Marienfeld, Germany, High-Precision 0117650) that were first sonicated in ethanol for 10 min, plasma-cleaned, then stored in isopropanol until use. U2OS cells were grown directly on the coverglasses in regular culture medium. The medium was changed after dye labeling and immediately before imaging into phenol red-free medium to reduce background, while all other components of the medium remained unchanged.

### Nucleofection

Because lipofection-based transfection methods often produce substantial background labeling in experiments with fluorescent dyes, for all imaging experiments involving exogenous expression we used the Lonza Amaxa II Nucleofector System with Cell Line Nucleofector Kit V reagent (Lonza VCA-1003). Briefly, U2OS cells were grown in 10 cm plates (ThermoFisher) for two days prior to nucleofection, trypsinized, spun down at 1200 rpm for 5 min, combined with vector and Kit V reagent according to manufacturer’s instructions, and nucleofected with program X-001 on an Lonza Amaxa II Nucleofector. After nucleofection, cells were immediately resuspended in regular culture medium at 37° C and plated onto coverslips. In all imaging experiments involving nucleofection, imaging was performed within 24 hours of plating.

### CRISPR/Cas9-mediated gene editing

Endogenous tagging of RARA in U2OS cells was performed with a protocol roughly following [16] with some modifications. This protocol relies on FACS sorting for cells that have been correctly modified to express HaloTag fused to the target protein. For U2OS cells, we nucleofected cells with plasmid expressing 3xFLAG-SV40NLS-pSpCas9 from a CBh promoter [58], mVenus from a PGK promoter, and guide RNA from a U6 promoter (pU6_sgRNA_CBh_Cas9_PGK_Venus_anti-RARA-C_terminus_1 and pU6_sgRNA_CBh_Cas9_PGK_Venus_anti-RARA-C_terminus_2), along with a second plasmid encoding the homology repair donor (pUC57_hom-Rep_RARA-HaloTag). The homology repair donor was built in a pUC57 backbone modified to contain HaloTag-3xFLAG with ∼500 base pairs of homologous genomic sequence on either side. Synonymous mutations were introduced at the cut site to prevent retargeting by Cas9. Each of the two guide RNA plasmids were nucleofected into separate populations of cells to be pooled for subsequent analysis. 24 hours after the initial nucleofection, we screened for mVenus-expressing cells using FACS and pooled these mVenus-positive cells in 10 cm plates. 5 days after plating, we labeled cells with HTL-TMR (Promega G8251) and screened for TMR-positive, mVenus-negative cells. Cells were diluted to single clones and plated in 96-well plates for a 2-3 week outgrowth step, during which the medium was replaced every 3 days. The 96-well plates were then screened for wells containing single colonies of U2OS cells, which were split by manual passage into two replicate wells in separate 96-well plates. One of these replicates was used to subpassage, while the other was used to harvest genomic DNA for PCR and sequencing-based screening for the correct homology repair product. In PCR screens, we used three primer sets: (A) primers external to the homology repair arms, expected to amplify both the wildtype allele and the edited allele (“PCR1”), (B) a primer internal to HaloTag and another external to it on the 5’ side, expected to amplify only the edited allele (“PCR2”), and (C) a primer internal to HaloTag and another external to it on the 3’ side, expected to amplify only the edited allele (“PCR3”). The primer sets for each target were the following. For RARA-GDGAGLIN-HaloTag-3xFLAG, we used prAH586 and prAH761 for PCR1, prAH761 and prAH762 for PCR2, and prAH763 and prAH764 for PCR3. For NPM1-GDGAGLIN-HaloTag-3xFLAG, we used prAH1092 and prAH1093 for PCR1, prAH1093 and prAH377 for PCR2, and prAH1092 and prAH373 for PCR3. U2OS cDNA from selected clones was isolated with DirectPCR Lysis Reagent (Viagen 101-T), treated with 0.5 mg/ml proteinase K for 15 min, incubated at 95° C for 1 hour, then subjected to PCRs 1 through 3 using Phusion polymerase in the presence of 5% DMSO. Amplicons from candidate clones were gel-purified (Qiagen 28704) and Sanger sequenced; only clones with the correct target sequence were kept for continued screening. A subset of these clones were chosen for characterization by Western blot, imaging, and luciferase assays.

For NPM1-GDGAGLIN-HaloTag-3xFLAG knock-in cell lines, we used a different strategy relying on nucleofected *S. pyogenes* Cas9 sgRNPs and linear dsDNA homology repair donors. The target insert (GDGAGLIN-HaloTag-3xFLAG from the vector PB PGKp-PuroR L30p MCS-GDGAGLIN-HaloTag-3xFLAG) was first amplified with ultramers encoding 120 bp homology arms (prAH867 and prAH868) using KAPA2G Robust HotStart polymerase (Kapa Biosystems KR0379) for 12 cycles. A small volume of this reaction was then used to seed a PCR reaction using primers prAH869 and prAH870 in Q5 High-Fidelity 2X Master Mix (Qiagen M0492). Products were purified by RNAClean XP magnetic beads (Beckman-Coulter A63987) and further cleaned by ethanol precipitation, followed by resuspension in a small volume of RNase-free water. For guides, we performed a three-primer PCR using prAH2000 and prAH2001 along with a unique oligo encoding the spacer (either prAH979 or prAH980) to produce a linear dsDNA product encoding the sgRNA preceded by a T7 promoter. We then used T7 RNA polymerase (NEB E2040S) to transcribe sgRNA from this template and purified the sgRNA with RNAClean XP magnetic beads according to manufacturer’s instructions. To assemble the sgRNP, we incubated 80 pmol sgRNA with 40 pmol purified SpyCas9-NLS (UC Berkeley Macrolab) for 15 min at 37° C in 20 mM HEPES pH 7.5, 150 mM KCl, 10 mM MgCl_2_, and 5% glycerol. sgRNPs were subsequent kept on ice and combined with donor immediately before nucleofection. For each nucleofection, we used 40 pmol sgRNP and 5 pmol dsDNA donor template suspended in <10 *µ*L with Lonza Amaxa Nucleofector II protocol X-001 in Lonza Kit V reagent. Roughly 1 million cells were used for nucleofection. Sorting for labeled cells, subcloning, and genotyping proceeded as previously described for RARA-GDGAGLIN-HaloTag-3xFLAG.

### Western blots

Antibodies were as follows. The ratio indicate the dilution factors used for Western blot. human TBP, Abcam Ab51841, 1:2500 (mouse); FLAG, Sigma-Aldrich F3165, 1:2000 (mouse).

For Western blots, cells were collected by scraping from plates in ice-cold PBS, then pelleted. Cell pellets were resuspended in lysis buffer (0.15 M NaCl, 1% NP-40, 50 mM Tris-HCl (pH 8.0), and a cocktail of protease inhibitors (Sigma-Aldrich 11697498001 dissolved in PBS with supplemented PMSF, aprotinin, and benzamidine), agitated for 30 min at 4° C, then centrifuged for 20 min at 12000 rpm, 4° C. The supernatant was then mixed with 2x Laemmli (to final 1x), boiled for 5 min, then run on 12.5% SDS-PAGE. After transfer to nitrocellulose, the membrane was blocked with 10% condensed milk in TBST (500 mM NaCl, 10 mM Tris-HCl (pH 7.4), 0.1% Tween-20) for one hour at room temperature. Antibodies were suspended in 5% condensed milk in TBST at the dilutions indicated above and incubated, rocking at 4° C overnight. After primary hybridization, the membrane was washed three times for 10 min with TBST at room temperature, hybridized with an anti-mouse HRP secondary antibody in 5% condensed milk in TBST for 60 min at room temperature, washed three more times with TBST for 10 min, then visualized with Western Lightning Plus-ECL reagent (PerkinElmer NEL103001) according to manufacturer instructions and imaged on a Bio-Rad ChemiDoc imaging system. Different exposure times were used for each antibody.

### Luciferase assays

All luciferase assays used pGL3-RARE-luciferase, a reporter containing firefly luciferase driven by an SV40 promoter with three retinoic acid response elements (RAREs). pGL3-RARE-luciferase was a gift from T. Michael Underhill (Addgene plasmid 13458; http://n2t.net/addgene:13458; RRID:Ad-dgene_13458) [57]. Luciferase assays were performed on cells cultivated in 6-well plates. Cells were transfected with 100 ng pGL3-RARE-luciferase and 10 ng pRL Renilla (Promega E2261) using Mirus TransIT-2020 Transfection Reagent (Mirus MIR 5404) for U2OS cells or Lipofectamine 3000 (ThermoFisher L3000015) for mES cells. Transfection was performed one day before assaying luciferase expression with the Dual-Luciferase Reporter Assay System (Promega E1910) according to manufacturer’s instructions. Readout was performed on a GloMax luminometer (Promega).

### Cell labeling

For spaSPT experiments, cells were labeled with one of two methods, depending on the type of dye. For non-photoactivatable fluorescent dyes including TMR-HTL (tetramethylrhodamine-HaloTag ligand; Promega G8251), we stained cells with 100 nM dye in regular culture medium for 10 min, then performed three 10 min incubations in dye-free culture medium separated by PBS washes. All PBS and culture medium was incubated at 37° C between medium changes and washes.

For experiments with photoactivatable dyes, which have lower cell permeability and slower wash in/wash out kinetics, we labeled cells with 100 nM dye in regular culture medium for 30 min, followed by four 30 min incubations in dye-free culture medium at 37° C. Between each incubation, we washed twice with PBS at 37° C. After the final incubation, cells were changed into phenol red-free medium for imaging.

### spaSPT

spaSPT experiments were performed with a custom-built Nikon TI microscope equipped with a 100X/NA 1.49 oil-immersion TIRF objective (Nikon apochromat CFI Apo TIRF 100X Oil), an EMCCD camera (Andor iXon Ultra 897), a perfect focus system to account for axial drift, an incubation chamber maintaining a humidified 37° C atmosphere with 5% CO_2_, and a laser launch with 405 nm (140 mW, OBIS, Coherent), 488 nm, 561 nm, and 633 nm (all 1 W, Genesis Coherent) laser lines. Laser intensities were controlled by an acousto-optic Tunable Filter (AA Opto-Electronic, AOTFnC-VIS-TN) and triggered with the camera TTL exposure output signal. Lasers were directed to the microscope by an optical fiber, reflected using a multi-band dichroic (405 nm/488 nm/561 nm/633 nm quad-band, Semrock) and focused in the back focal plane of the objective. The angle of incident laser was adjusted for highly inclined laminated optical sheet (HiLo) conditions [12]. Emission light was filtered using single band-pass filters (Semrock 593/40 nm for PAJFX549 and Semrock 676/37 nm for PAJF646). Hardware was controlled with the Nikon NIS-Elements software.

For stroboscopic illumination, the excitation laser (561 nm or 633 nm) was pulsed for 1-2 ms (most commonly 1 ms) at maximum (1 W) power at the beginning of the frame interval, while the photoactivation laser (405 nm) was pulsed during the ∼447 *µ*s camera transition time, so that the background contribution from the photoactivation laser is not integrated. For all spaSPT, we used an EMCCD vertical shift speed of 0.9 *µ*s and conversion gain setting On our setup, the pixel size after magnification is 160 nm and the photon-to-grayscale gain is 109. 15000-30000 frames with this sequence were collected per nucleus, during which the 405 nm intensity was manually tuned to maintain low density of fluorescent particles per frame.

### Localization and tracking

To produce trajectories from raw spaSPT movies, we used a custom tracking tool publicly available on GitHub (`quot`, available at https://github.com/alecheckert/quot) that provides a graphical user interface for comparing detection, subpixel localization, and tracking algorithms adapted from other sources. All localization and tracking for this manuscript was performed with the following settings:

- *Detection*: generalized log likelihood ratio test with a 2D Gaussian kernel of fixed radius 190 nm (detection method `llr` with `k` = 1.2, a 15 pixel window size (`w` = 15), and a log ratio threshold of 16.0 (`t` = 16.0; inspired by [59]).
- *Subpixel localization*: Levenberg-Marquardt fitting of a 2D integrated Gaussian point spread function model (quot localization method `ls_int_gaussian`; inspired by [61]) with fixed radius 190 nm, window size 9 pixels, maximum 20 iterations per PSF, with a damping term of 0.3 for parameter updates. The 2D integrated Gaussian PSF model is described, for instance, in [62]. We used the radial symmetry method [60] to make the initial guess used to start the Levenberg-Marquardt algorithm.
- *Tracking*: quot tracking algorithm `conservative` with a 1.2 *µ*m search radius. This simple algorithm searches for particle-particle reconnections that are “unambiguous” in the sense that no other reconnections are possible within the specified search radius. These reconnections are then used to synthesize trajectories. At high densities, many jumps are discarded because other reconnection possibilities given the search radius exist.

After localization and tracking, all trajectories in the first 1000 frames of each movie were discarded. Localization density tends to be high in these frames, so they can contribute tracking errors that compromise accuracy. The mean localization density for most movies in the remaining set of frames was less than one emitter per frame.

For experiments involving HaloTag or HaloTag-NLS, which move quickly, we used a broader search radius at 2.5 *µ*m. All other settings were kept the same.

### Spinning disk confocal imaging

Experiments using spinning disk confocal imaging were performed at the UC Berkeley High-Throughput Screening Facility on a Perkin Elmer Opera Phenix equipped with a controller for 37° C and 5% CO_2_, using a built-in 40X water immersion objective.

### Simulation

All simulations were performed with a simple publicly available spaSPT simulation tool (strobesim; https://github.com/alecheckert/strobesim). This tool generates trajectories for a variety of different motion types - for instance, Brownian motion, fractional Brownian motion, or Levy flights - or accepts a user-defined type of motion, and simulates the act of observing this motion in a thin focal plane in a particular cellular geometry. The tool provides ways to set the photobleaching and geometry settings for the spaSPT simulation.

Unless otherwise noted, in this manuscript we used a spherical cell geometry with radius 5 *µ*m and a focal plane with 700 nm depth bisecting the sphere. Simulated emitters were subject to photoactivation and photobleaching throughout the sphere and were only observed when their positions coincided with the focal volume. We simulated sparse tracking without gaps, so that if an emitter passed twice through the focal volume, it counted as two separate trajectories. At the sparsity used for these simulations, tracking is unambiguous and so tracking errors do not contribute to the outcome.

For discrete-state simulations, the number of particles in each state was modeled as a multinomial random variable drawn from the underlying state occupancies. As a result, there is an inherent variability associated with the “true” fractional occupancies for each simulation replicate, exactly as would be expected in spaSPT experiments.

For simulations with state transitions, we modeled the particles as two-state Markov chains with identical transition rates between the states. Each state was associated with a constant diffusion coefficient. These Markov chains were simulated on subframes grained at 100 iterations per frame interval. For instance, for simulations with 7.48 ms frame intervals, the underlying Markov chain was simulated on subframes of 74.8 *µ*s. During each subframe, the state of the MC was assumed to be constant and we simulated diffusion according to the Euler-Maruyama scheme with the current diffusion coefficient. The positions of the particle at the frame interval were recorded.

### DPMM and SA implementation

The gamma approximation DPMM as described in the Supplementary Information has a publicly available implementation (dpsp; https://github.com/alecheckert/dpsp). This implementation is a Python interface to an underlying C++ algorithm for Gibbs sampling from the DPMM posterior distribution. Some examples are provided in the repository.

The state array (SA) as described in the Supplementary Information has a publicly available pure Python implementation (spagl; https://github.com/alecheckert/spagl). This repository also provides a variety of simple interfaces to produce the plots shown in the manuscript, including the prior/posterior diffusion coefficient likelihoods as a function of space (Figure 5B) and as a function of time (Figure 5D).

## Figures

**Supplementary Movie 1: Example of spaSPT data.** NPM1-HaloTag in U2OS osteosarcoma nuclei was labeled with 100 nM PA-JFX549-HTL for 5 min followed by washes as described in Materials and Methods, then imaged with a HiLo setup at 7.48 ms frame intervals with 1.5 ms excitation pulses. The pixel size after accounting for magnification is 160 nm. Dots and lines indicate the output of the detection and tracking algorithm; each trajectory has been given a distinct color.

**Supplementary Movie 2: Illustration of defocalization for a single regular Brownian state**. Trajectories were simulated in a 5×5×10 *µ*m ellipsoid *µ*m using the Euler-Maruyama scheme for regular Brownian motions with specular reflections at the ellipsoid boundaries. The diffusion coefficient for all trajectories was held constant at 2.0 *µ*m^2^ s^*−*1^, while trajectories were randomly photoactivated at any point in the sphere and were subject to Poisson bleaching at 14 Hz. The left panel shows the three-dimensional context of the trajectories, with dotted lines indicating the boundaries of the focal volume. The depth of the focal volume was 700 nm, which is roughly equivalent to the measured depth of field for our oil immersion objectives. The right panel shows the projection of the trajectories that coincide with the focal volume onto a hypothetical camera. Notice that particles may make multiple transits through the focal volume that manifest as distinct trajectories.

**Supplementary Movie 3: Illustration of defocalization for multistate regular Brownian motion**. Trajectories were drawn from two states - a fast state with diffusion coefficient 5.0 *µ*m^2^ and a slow state with diffusion coefficient 0.05 *µ*m^2^ s^*−*1^ - and simulated with a spherical nucleus with 5 *µ*m radius. Similar to Supplementary Movie 2, the left panel shows the trajectories in their native three dimensions while the right panel shows trajectories as projected through the focal volume onto the surface of a camera.

**Figure S1:**
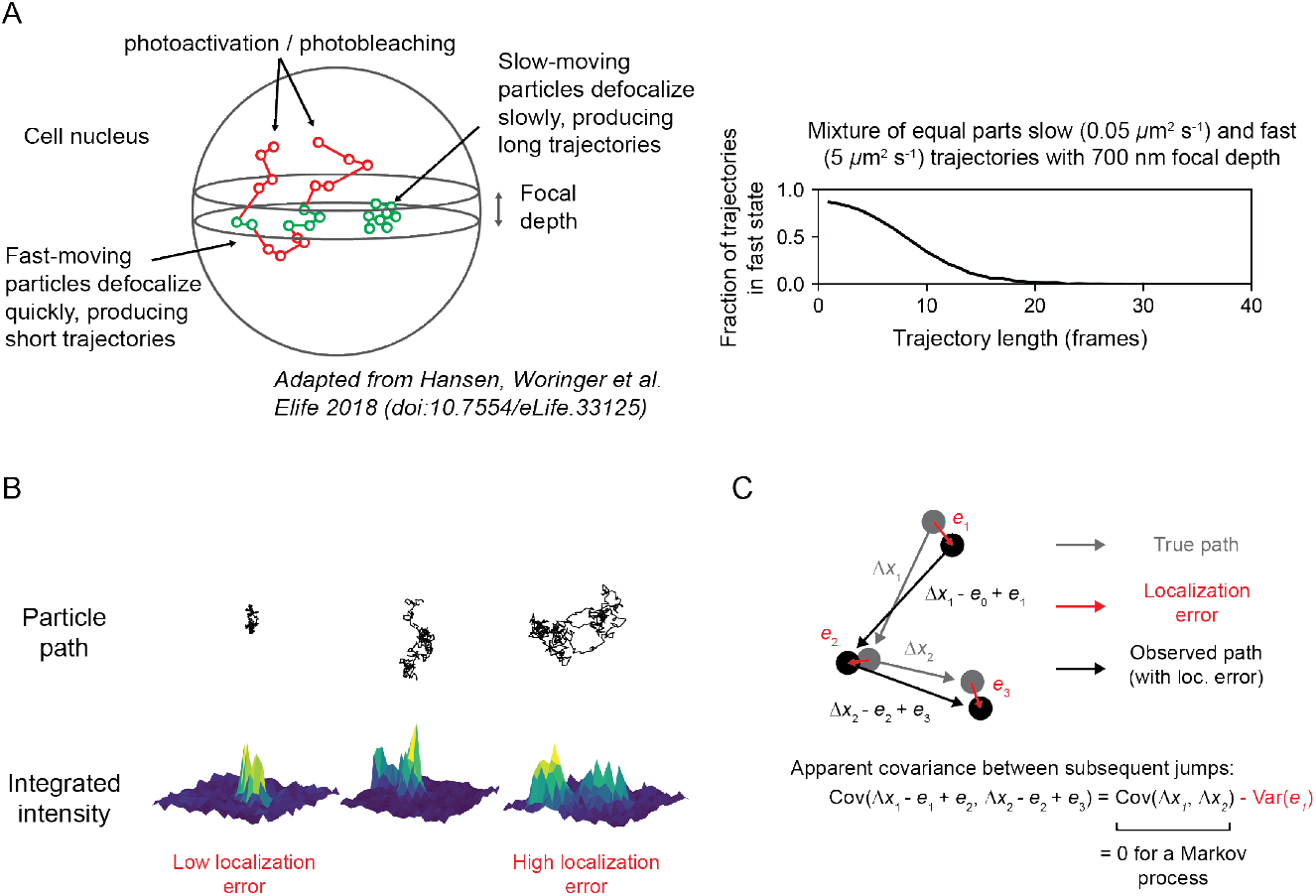
Challenges with using the trajectory-averaged mean-squared displacement (MSD) for multiple diffusing states in a thin focal volume. (A) Schematic of the state bias issue arising from thin focal volumes. For the simulation, 340727 trajectories were drawn with equal probability from a slow state (0.001 *µ*m^2^ s^*−*1^) and a fast state (5 *µ*m^2^ s^*−*1^) and were simulated in a 5 *µ*m radius nucleus with a 700 nm focal depth and 14 Hz bleaching rate. For each trajectory length, the fraction of trajectories in the fast state was quantified. (B) Illustration of the relationship between mobility and localization error. Fast-moving molecules present blurrier spots, leading to higher localization error. (C) Illustration of the role of localization error in the jump variance and covariance between subsequent jumps. As a result, even Markov processes such as regular Brownian motion have nonzero covariance between subsequent jumps when measured with localization error.

**Figure S2:**
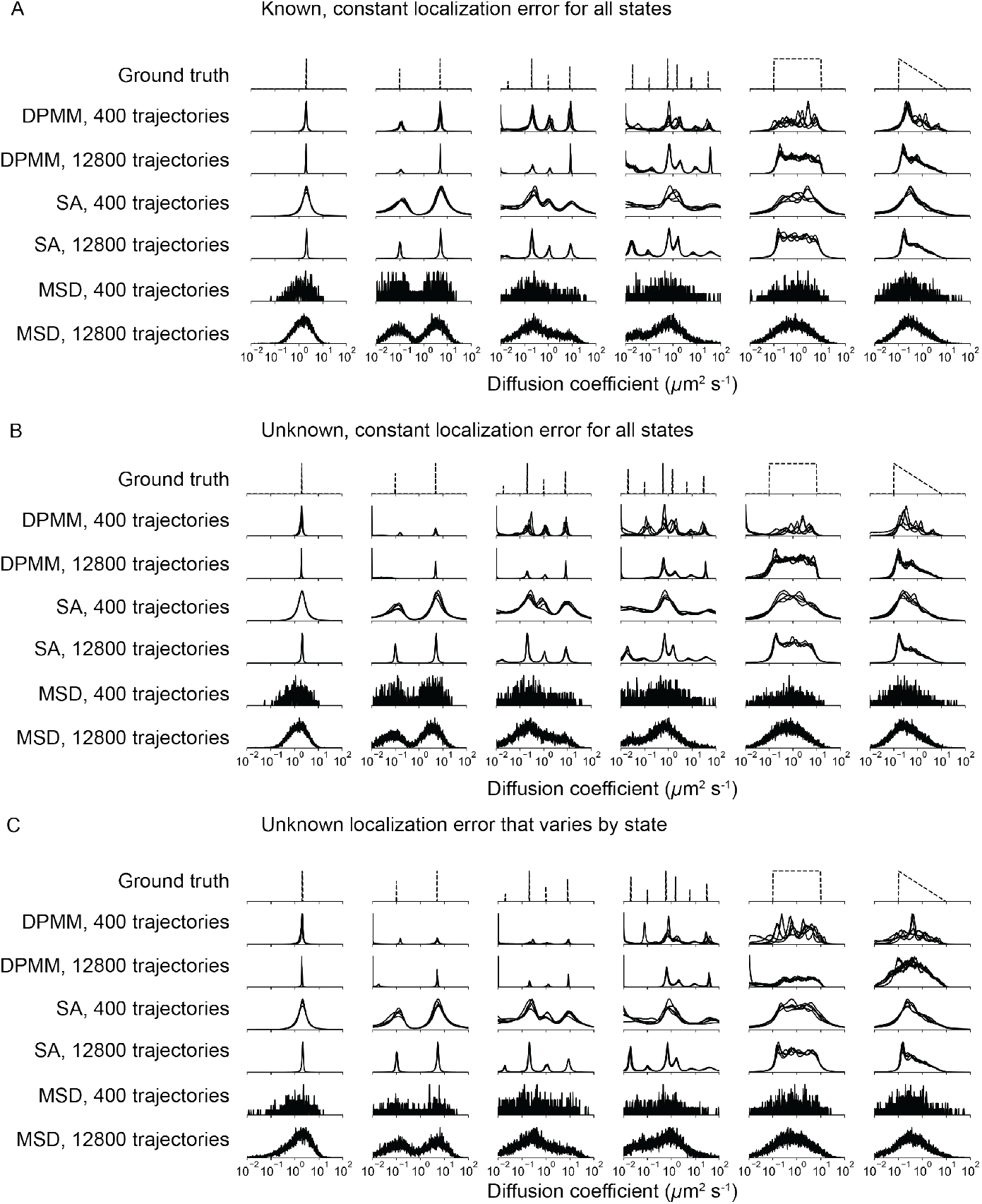
Comparison of the MSD, DPMM, and SA approaches on simulated mixtures of regular Brownian motions with error. In all panels, the y-axis represents the probability (or inferred probability) of a particular diffusion coefficient. For the DPMM and SA approaches, this is inferred as the mean or marginal mean of the posterior distribution, respectively. In the case of the MSD approach, we instead report the proportion of trajectories that fall into the corresponding bin. For the SA method, we used a parameter grid with diffusion coefficients spanning 0.01 - 100.0 *µ*m^2^ s^*−*1^ and localization errors spanning 0 - 60 nm. Five independent replicates are overlaid on each subplot. (A) Simulations with known and constant localization error (standard deviation 30 nm). (B) Simulations with constant but unknown localization error (standard deviation 30 nm). In the cases of DPMM and MSD, the localization error was first inferred using jump covariance and then held constant when estimating the distribution over the diffusion coefficient. In the case of SA, the localization error is jointly inferred with the diffusion coefficient. (C) Simulations with variable and unknown localization error. Inference proceeded as in (B), but in the case of the MSD approach, the localization error and diffusion coefficient were jointly inferred for each trajectory. The localization errors for each simulation are recorded in the Supplementary Methods.

**Figure S3:**
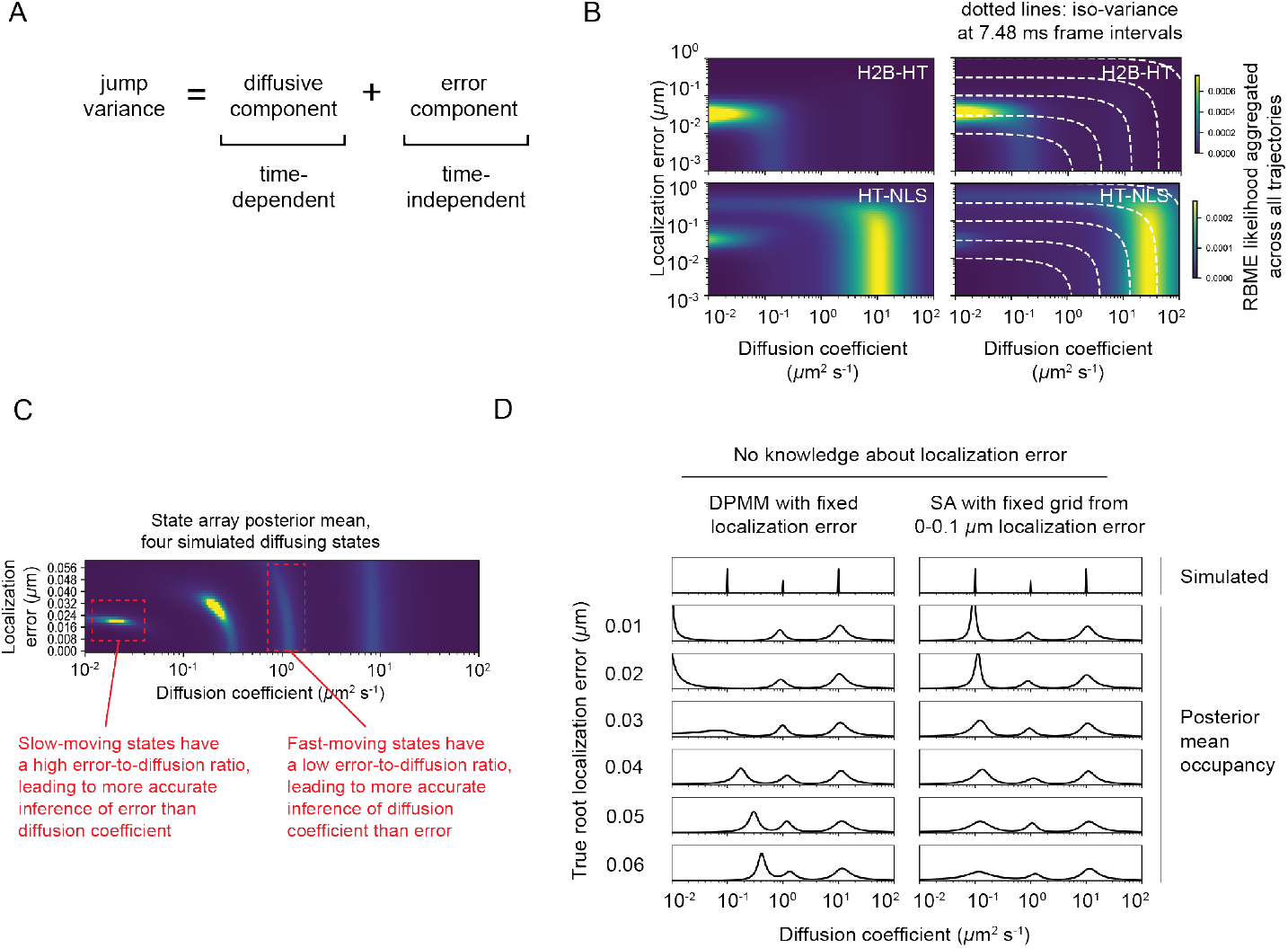
Illustration of the connection between diffusion coefficient and localization error. (A) The variance of the jumps taken by an RBME is the sum of contributions from diffusion and localization error. These contributions are distinguishable when considering multiple frame intervals because the diffusive component scales with time, whereas the error component does not. However, estimating either the error or diffusive component is more difficult when the component is small compared to the other. (B) Aggregated RBME likelihoods for two experimental spaSPT datasets. Dotted white lines indicate contours with constant jump variance. (C) Illustration of the difficulty inherent in distinguishing the effects of diffusion and error for slow or fast diffusion coefficients for a simulated spaSPT dataset with four states. (D) Illustration of the danger of misestimating the localization error for the DPMM method with simulated mixtures of three RBME states. In this case, the assumed localization error in the DPMM algorithm was held constant at 30 nm. Because SAs naturally incorporate uncertainty about localization error, inference is more stable with respect to changes in the experimental localization error.

**Figure S4:**
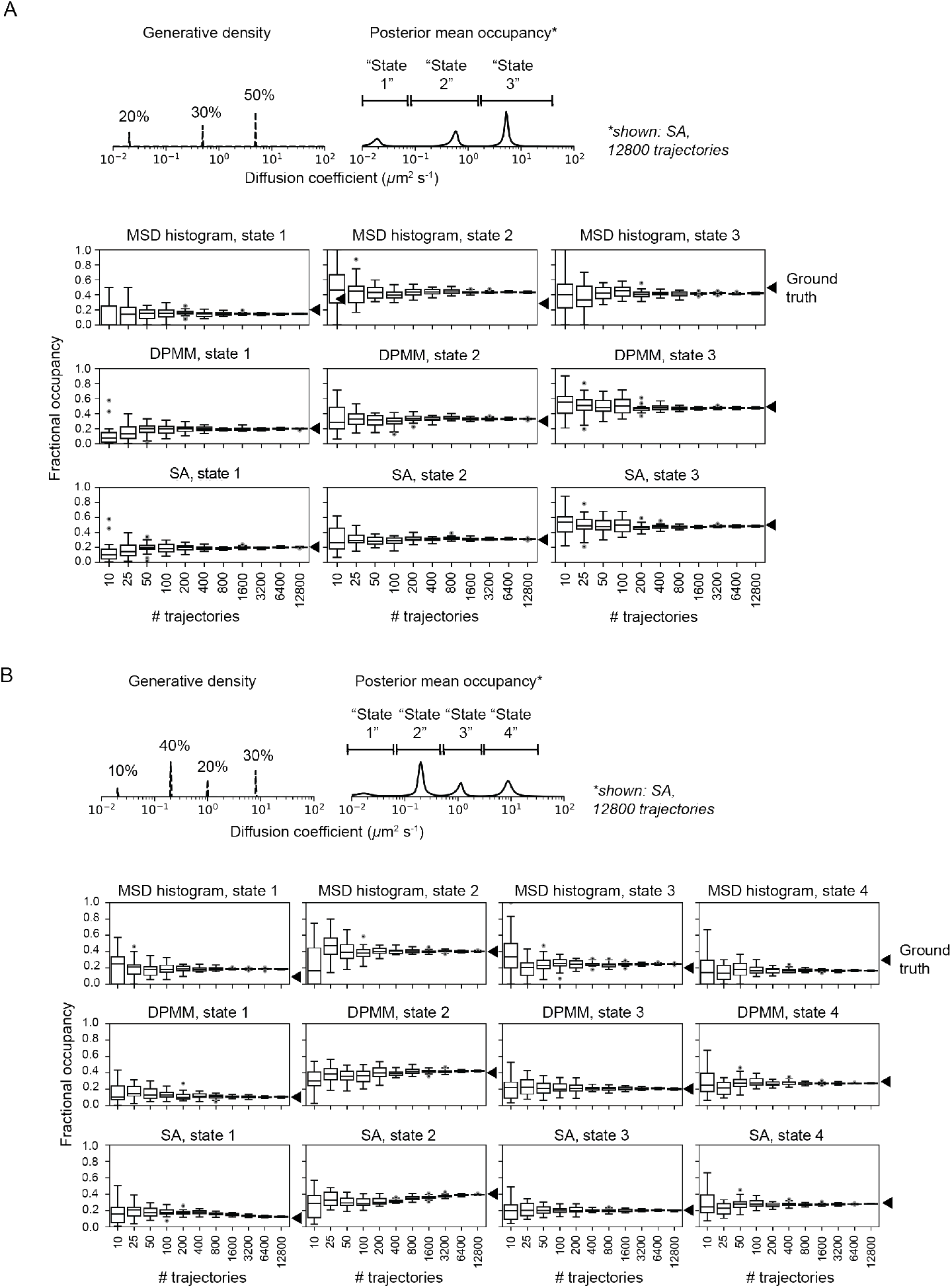
Comparison of state occupation accuracy for MSD, DPMM, and SA approaches. (A) Retrieving state occupations for a three-state model. Fractional occupation was estimated by integrating the normalized histogram (for the MSD method) or by integrating the mean of the posterior distribution (for DPMM and SA methods). The limits of integration for each state were determnined based on visual inspection of the posterior distribution. The limits of integration for each state were 0 - 0.08 *µ*m^2^ s^*−*1^ for the first state, 0.08 - 1.5 *µ*m^2^ s^*−*1^ for the second state, and 1.5 - 40 *µ*m^2^ s^*−*1^ for the third state. (B) Retrieving state occupations for a four-state model. Fractional occupation was estimated as in (A). The limits of integration for each state were 0 - 0.08 *µ*m^2^ s^*−*1^ for the first state, 0.08 - 0.5 *µ*m^2^ s^*−*1^ for the second state, 0.5 - 3 *µ*m^2^ s^*−*1^ for the third state, and 3 - 40 *µ*m^2^ s^*−*1^ for the fourth state.

**Figure S5:**
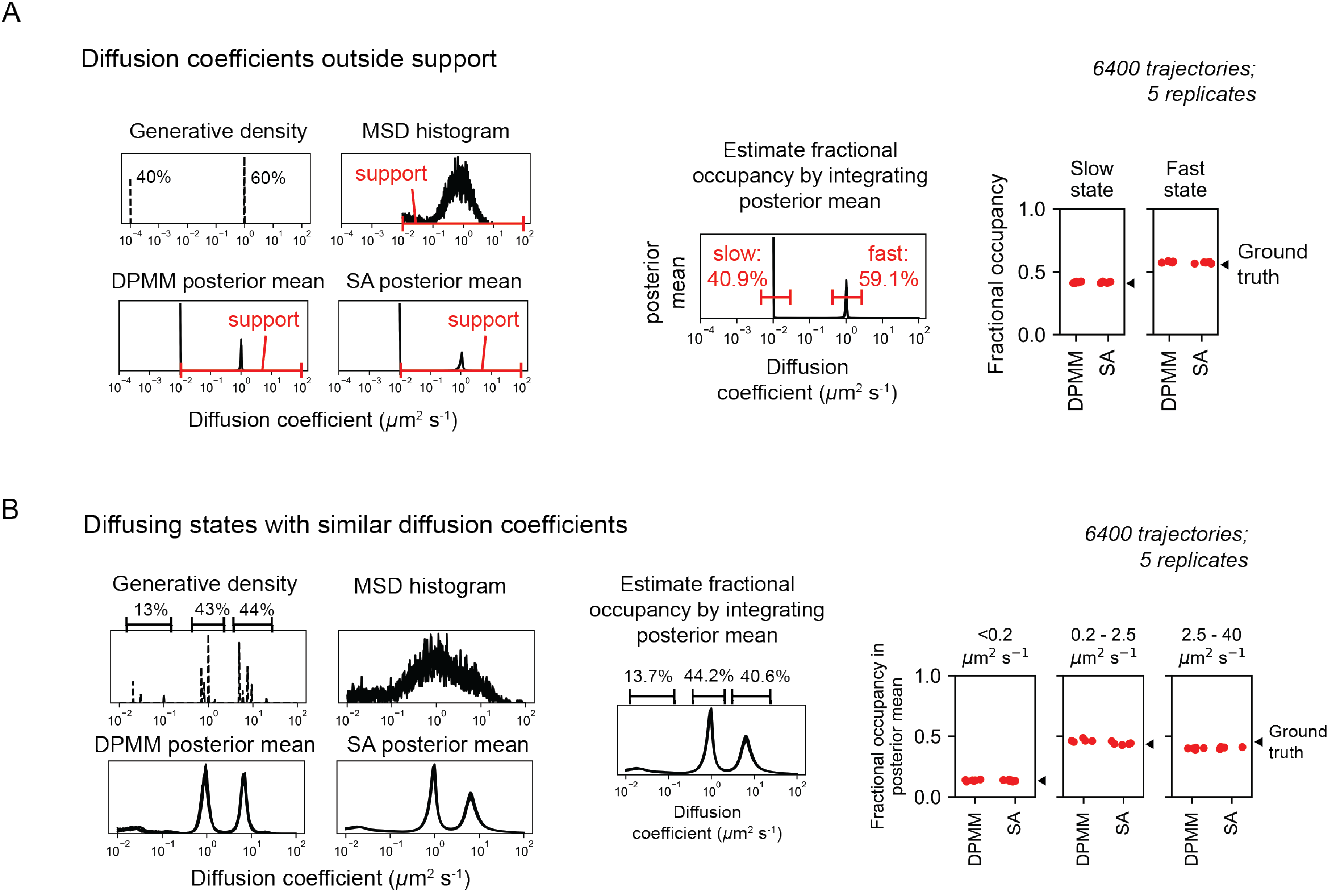
Comparison of the DPMM and SA algorithms on two difficult situations. (A) Running DPMM and SA on trajectories with diffusion coefficients slower than the minimum diffusion coefficient included in the support. Fractional occupation for each state was estimated by integrating the posterior mean over the peaks. For the MSD histogram, DPMM posterior mean, and SA posterior mean subplots, five independent replicates are overlaid onto the same plot. (B) Running DPMM and SA on diffusing states with similar diffusion coefficients. For the MSD histogram, DPMM posterior mean, and SA posterior mean subplots, five independent replicates are overlaid onto the same plot.

**Figure S6:**
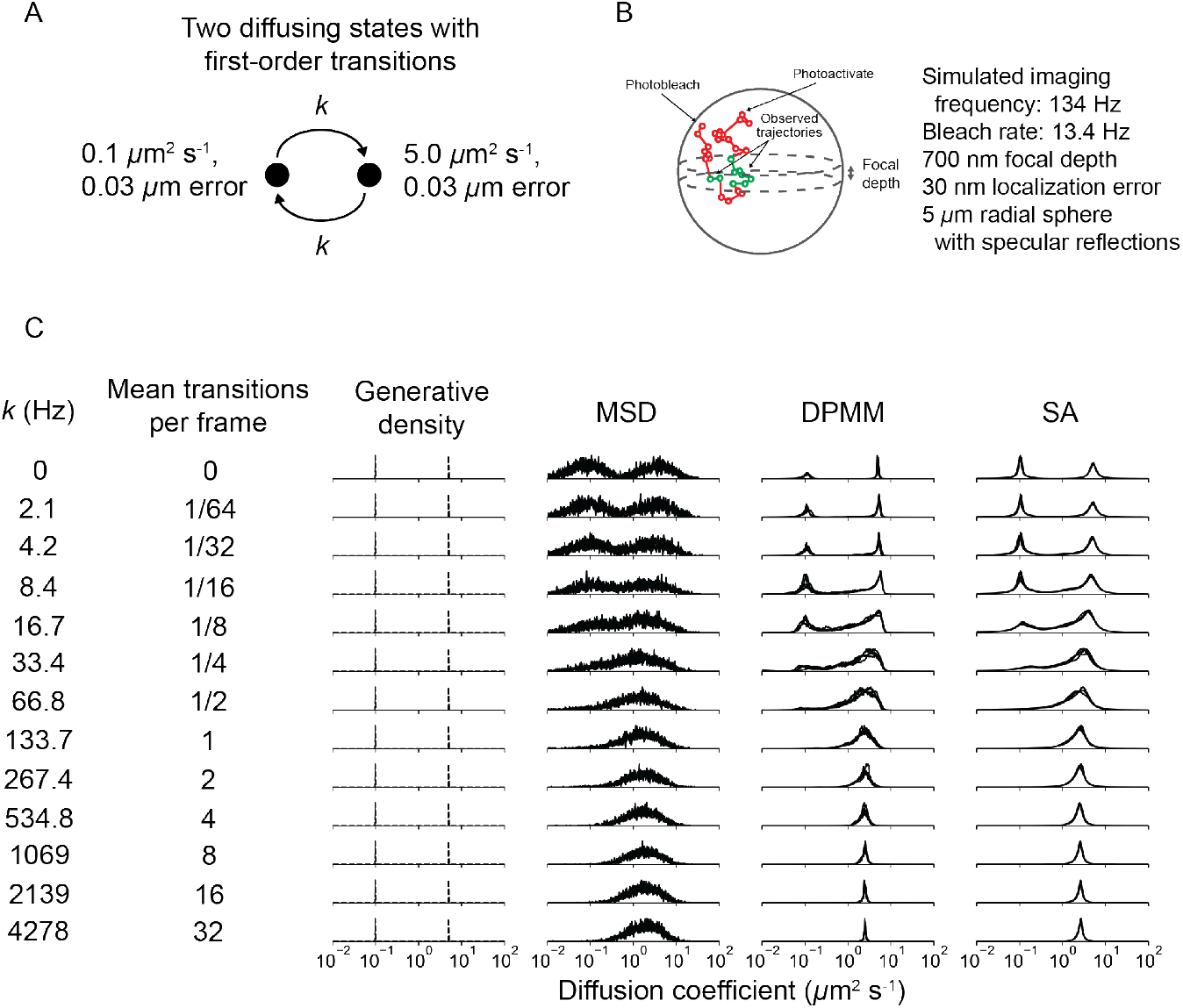
Effect of state transitions on the outcome of the DPMM and SA inference methods. (A) State diagram for two-state regular Brownian motion with state transitions. The transition rate constant *k* was identical for both transitions. (B) Settings for the state transition simulations. Under these conditions, the mean trajectory length was 7 frames. 6400 trajectories were used for each inference run. (C) Outcome of the transition simulations. The y-axis corresponds to state occupation. For both DPMMs and SAs, we used a maximum trajectory length of 12 frames. For the MSD, DPMM, and SA methods, the result of inference with five independent replicates are overlaid on each subplot.

**Figure S7:**
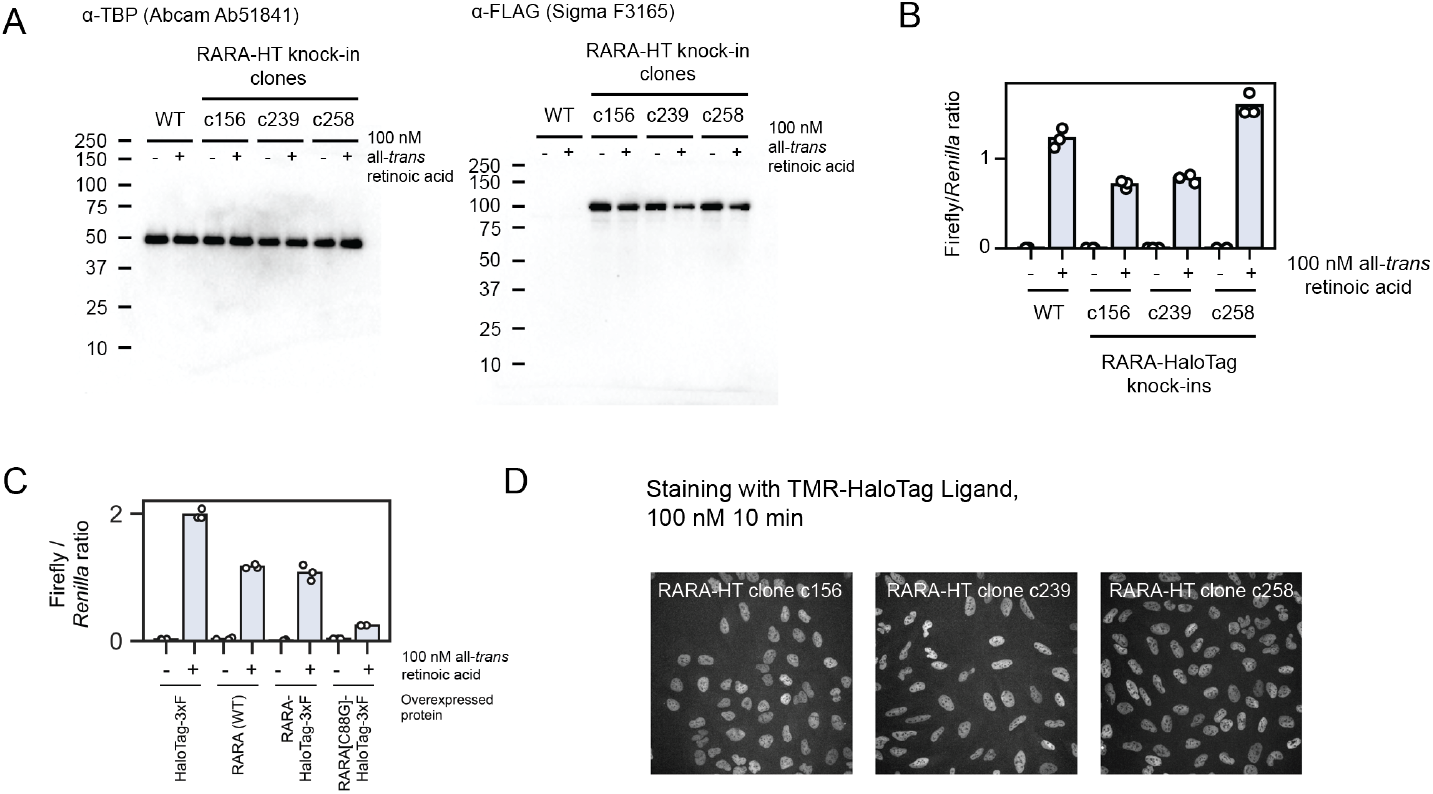
Validation of endogenously tagged U2OS RARA-HaloTag cell lines. (A) Western blots for endogenous RARA-HaloTag knock-ins in U2OS nuclei. The expected molecular weight of RARA-HaloTag-3xFLAG is 97 kDa. (B) Luciferase assays with a retinoic acid-responsive promoter with wildtype or endogenously tagged U2OS cell lines. (C) Luciferase assays with transfected RARA constructs to assess the effect of tagging on transactivation of a retinoic acid response element-driven luciferase gene. RARA(WT) indicates a transgene bearing the wildtype version of RARA, and C88G is a DNA-binding mutant. (D) Spinning disk confocal microscopy images of TMR-labeled endogenously tagged RARA-HaloTag cell lines. The intensities for all three images have been identically scaled.

**Figure S8:**
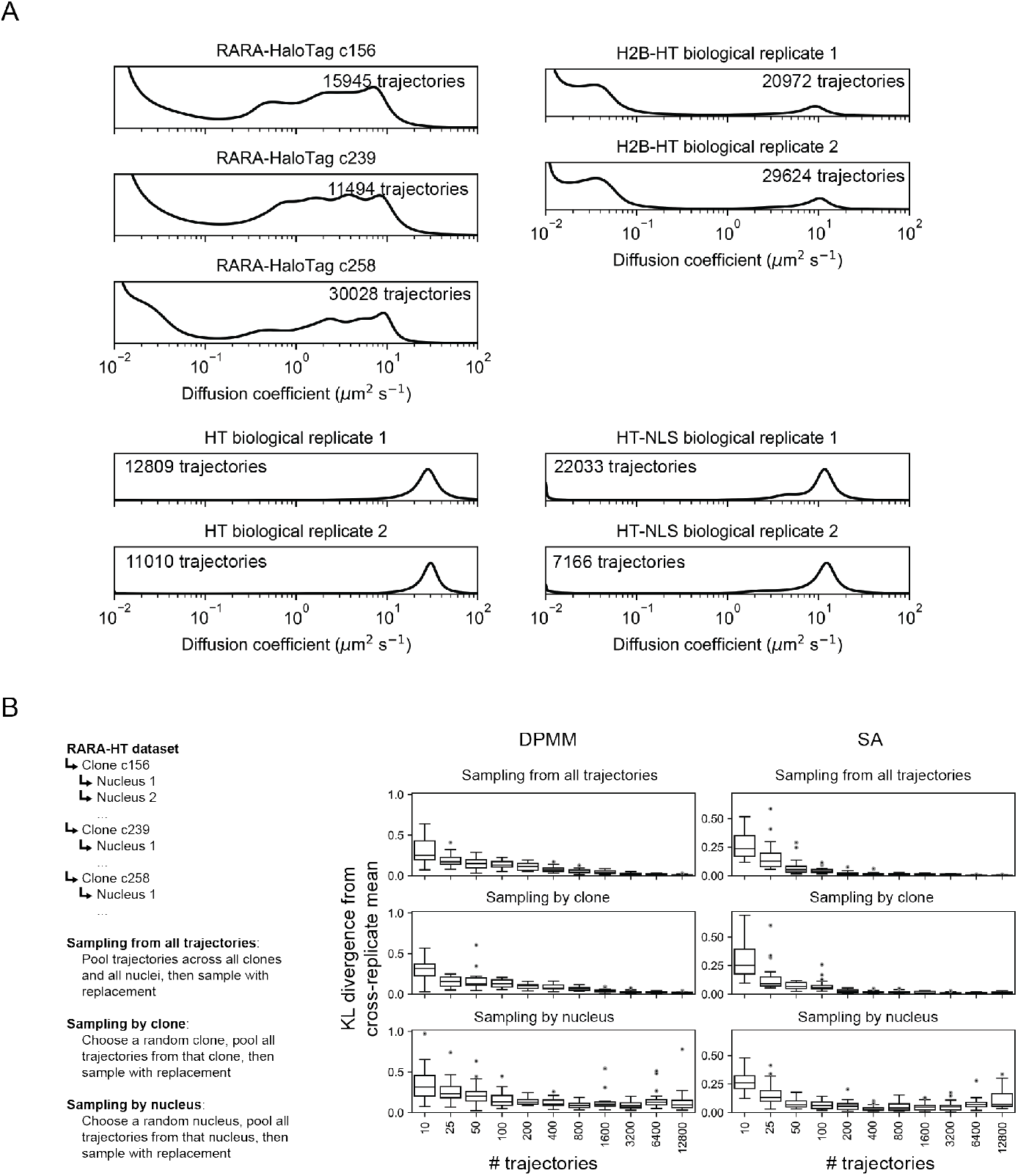
Assessing the variability of the SA posterior mean for experimental spaSPT datasets. (A) Biological replicates of each of the tracking experiments shown in Fig. 4A. Tracking was performed on an inverted TIRF setup with 7.48 ms frame intervals and 1 ms stroboscopic excitation with the dye PAJFX549-HaloTag ligand. RARA-HT cells were knock-ins as described in Fig. S7, H2B-HaloTag cells were previously described [52], and HT and HT-NLS were expressed from a nucleofected PiggyBac vector under an EF1*α* promoter. (B) Bootstrap aggregation to evaluate the sources of variability in DPMM/SA runs on RARA-HaloTag trajectories. RARA-HaloTag trajectories from one tracking dataset were sampled according to one of three schemes, then analyzed with the DPMM and SA methods. 20 replicates were performed for each condition, and the Kullback-Leibler divergence of the posterior mean of each replicate from the cross-replicate posterior mean were used to quantify variability.

**Figure S9:**
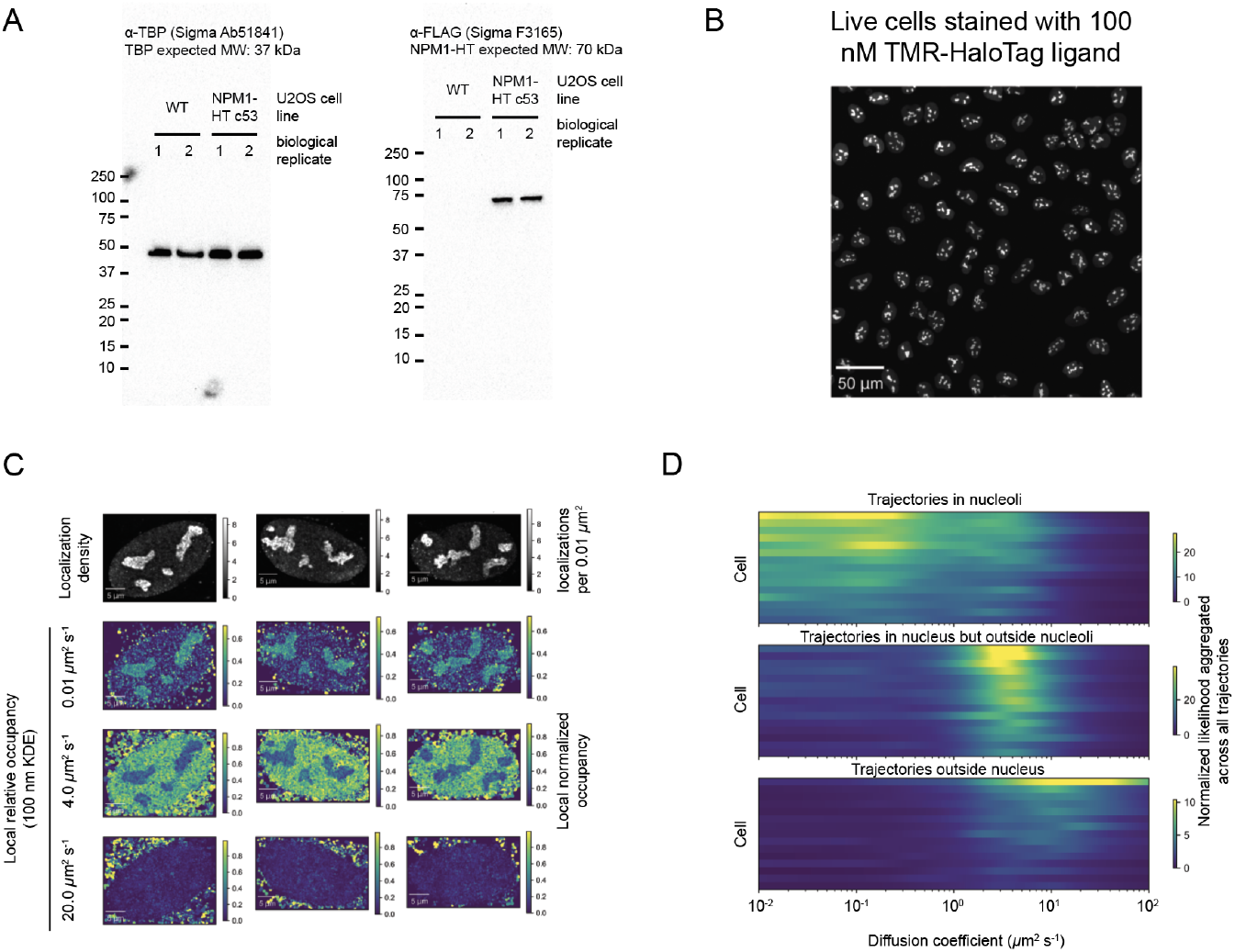
Supplementary data related to Figure 5. (A) Western blots on heterozygously tagged NPM1-HaloTag-3xFLAG in U2OS nuclei. (B) U2OS NPM1-HaloTag cells stained with 100 nM tetramethylrhodamine and imaged on a spinning disk confocal microscope. (C) Additional examples of cells quantified as in Fig. 5B. (D) Aggregate RBME likelihood functions for trajectories in different subcellular compartments. Trajectories were classified as either inside nucleoli, “nucleoplasmic” (outside nucleoli but inside the nucleus), or extranuclear. The RBME likelihood function was evaluated on each set of trajectories, aggregated across trajectories, and plotted as a function of the diffusion coefficient. Each row in the plot is separate nucleus, and likelihoods have been scaled for each compartment by the total number of trajectories.

**Figure S10:**
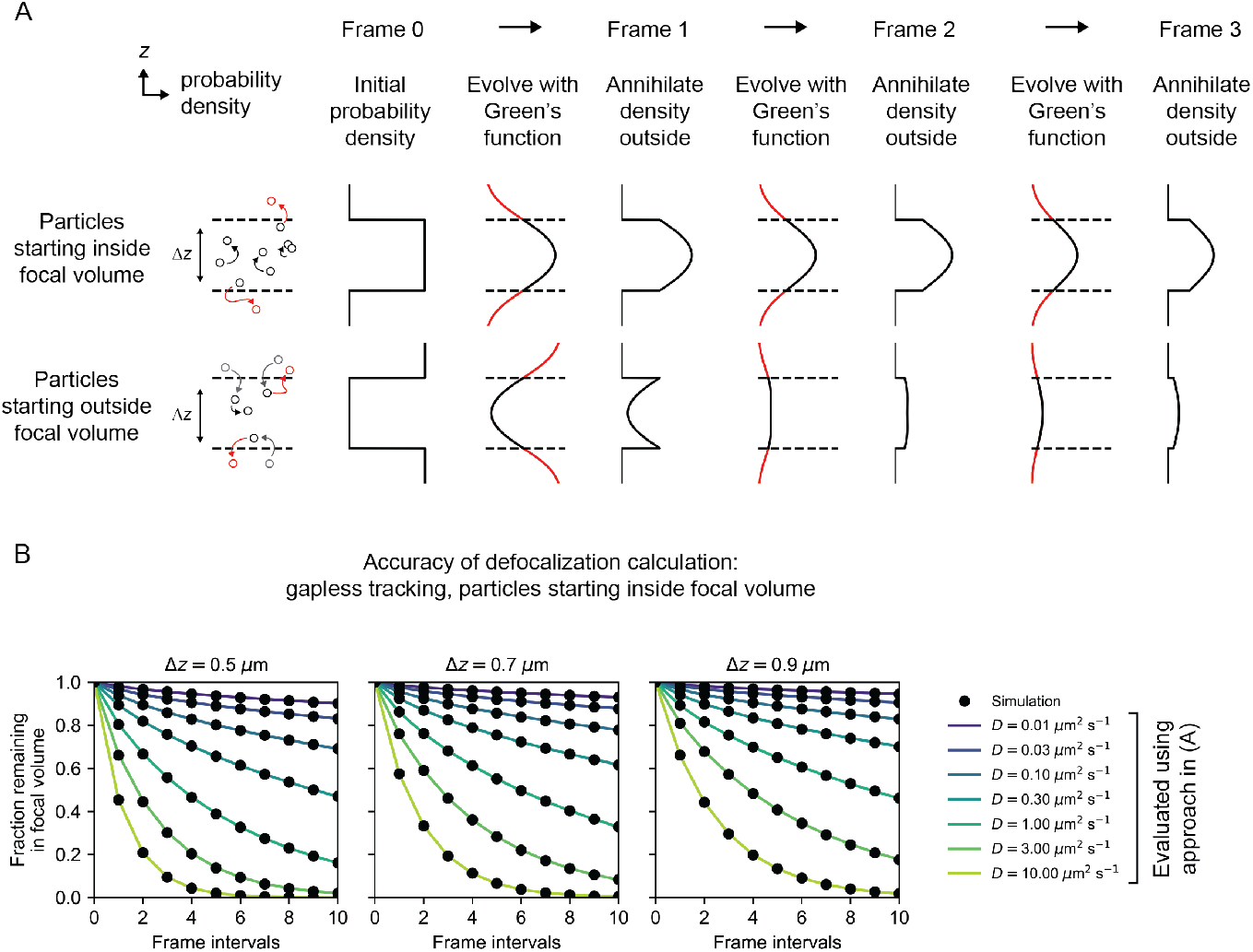
Approach to calculate the defocalized fraction for Markov processes tracked without gaps. (A) Schematic for the approach used to calculate the fraction of defocalized trajectories in 2D microscopy setups. Two possible initial probability densities for the particle in *z* are shown. The Green’s function for the diffusion model, estimated from motion in the XY plane, is used to propagate the probability density. At each frame interval, the density that lies outside the focal volume (corresponding to particles that are not observed) is set to zero. (B) Comparison of the algorithm in (A) with the defocalized fraction from simulated data. Trajectories were computationally photoactivated in a slab with thickness Δ*z* and infinite XY extent, then tracked without gaps. The fraction of trajectories remaining was quantified at each frame interval. Each black dot corresponds to a simulation with 100000 trajectories, while the lines correspond to the output of the algorithm in (A).

**Figure S11:**
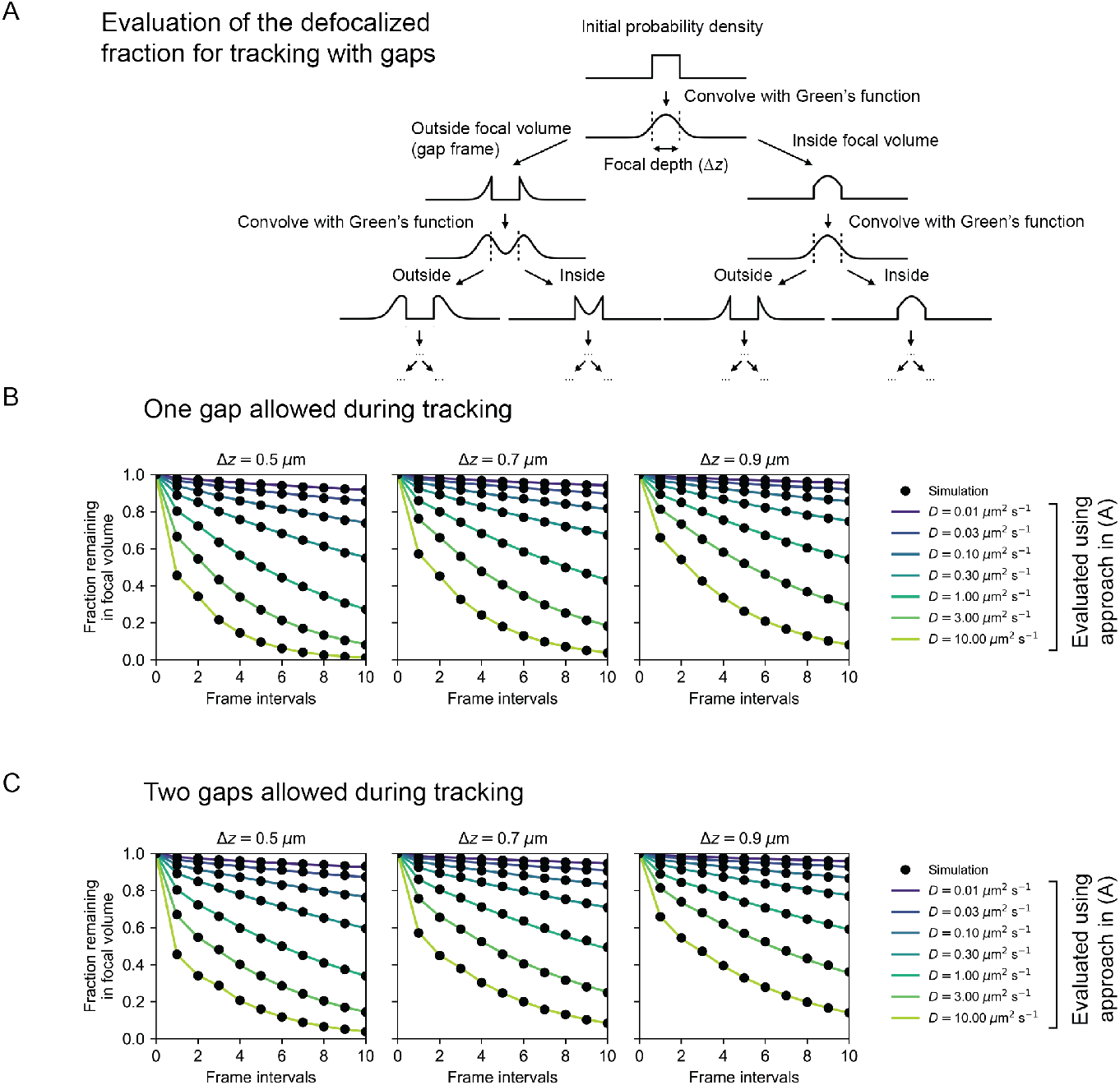
Approach to calculate the defocalized fraction for Markov processes tracked with gaps. (A) Schematic for the approach used to calculate the fraction of defocalized trajectories in 2D microscopy setups, tracked with gap frames allowed. Because tracking with *n* gaps allows particles to spend up to *n* frames outside the focal volume before returning, this approach alternates between propagating the probability density with the Green’s function for the Markov process and recursively partitioning the probability density into components inside and outside the focal volume, aggregating the density that lies outside the focal volume to calculate the defocalization function. An iterative version of the algorithm is outlined in the Supplementary Information. (B) Comparison of the algorithm in (A) with the defocalized fraction from simulated data with one gap allowed during tracking, as in Fig. **??**B. (C) Identical to (B), except two gaps were allowed during tracking.

## Notes

### Competing Interest Statement

The authors have declared no competing interest.

