## Supplementary Info for "Recovering mixtures of fast diffusing states from short single particle trajectories"

### Supplementary Information

**D Appendix: Subdiffusion with localization error** **26**

### 1 Overview

We describe two methods to infer mixtures of diffusive states from a collection of observed trajectories in an spaSPT experiment. Both approaches are rooted in Bayesian mixture models. In this introductory section, we describe the problem that both methods attempt to address and derive a mixture model appropriate for the fragmented trajectories collected in typical spaSPT experiments. The rest of the sections focus on methods to interpret the posterior distribution for this model.

#### 1.1 Statement of problem

We start with an observed sequence of $N$ trajectories $\mathbf{X} = (X_1, ..., X_N)$. We consider the $i^{\text{th}}$ trajectory $X_i$ as a random variable distributed according to a PDF $f_X(x|\theta_j)$, where $\theta_j$ is a vector of parameters governing its motion and $j \in \{1, 2, ..., K\}$ is the *state* of trajectory $i$. As an example, for regular Brownian motion, $\theta_j$ would be the diffusion coefficient and localization error associated with diffusive state $j$ (see Appendix A).

If we have $K$ states, then let $\tau_j$ be the fractional occupation of state $j$, so that $0 \leq \tau_j \leq 1$ and $\sum_{j=1}^{K} \tau_j = 1$. We use $\boldsymbol{\tau}$ to denote the vector of all state occupations and $\boldsymbol{\theta}$ to denote the matrix of all state parameters.

We assume that each trajectory is generated from one state. (In other words, we neglect state transitions.) Then we can represent the trajectory-state assignments as a $N$-by-$K$ matrix $\mathbf{Z}$, so that $Z_{ij} = 1$ if trajectory $i$ originates from state $j$ and $Z_{ij} = 0$ otherwise [10]. Each row, $Z_i$, is a vector with one element set to 1 and the other elements set to 0. This kind of definition makes it convenient to express the probability of $\mathbf{Z}$ given some state occupations $\boldsymbol{\tau}$ as

$$p(\mathbf{Z} \mid \boldsymbol{\tau}) = \prod_{i=1}^{N} \prod_{j=1}^{K} \tau_j^{Z_{ij}}$$

as well as the joint probability of $\mathbf{X}$ and $\mathbf{Z}$:

$$p(\mathbf{X}, \mathbf{Z} \mid \boldsymbol{\tau}, \boldsymbol{\theta}) = \prod_{i=1}^{N} \prod_{j=1}^{K} \left( \frac{\tau_j f_X(X_i|\theta_j)}{\sum_{k=1}^{K} \tau_k f_X(X_i|\theta_k)} \right)^{Z_{ij}}$$

In cellular spaSPT experiments, $\boldsymbol{\tau}$, $\boldsymbol{\theta}$, $K$, and $\mathbf{Z}$ are typically unknown. Our goal is to learn about these parameters given the observed trajectories. The Bayesian approach is to model all parameters as random variables and evaluate their conditional ("posterior") distribution given $\mathbf{X}$. Using Bayes' theorem, the posterior distribution can be expressed

$$p(\boldsymbol{\tau}, \boldsymbol{\theta}, K, \mathbf{Z} \mid \mathbf{X}) = \frac{p(\mathbf{X} \mid \boldsymbol{\tau}, \boldsymbol{\theta}, K, \mathbf{Z})\, p(\boldsymbol{\tau}, \boldsymbol{\theta}, K, \mathbf{Z})}{p(\mathbf{X})} \tag{1}$$

The term $p(\boldsymbol{\tau}, \boldsymbol{\theta}, K, \mathbf{Z})$, the probability of model parameters in the absence of observed trajectories, is the *prior distribution* and represents existing knowledge about the parameters before doing the experiment.

For example, if we have a known, fixed number of states $K$ and no reason *a priori* to favor one state over another, we choose a Dirichlet prior for $\boldsymbol{\tau}$ and some prior $H$ for each $\theta_j$. ($H$ is often chosen to be conjugate to $f_X(x|\theta)$ for mathematical simplicity.) Together this defines the finite-state mixture model (FSMM), a workhorse of stochastic modeling:

$$\begin{aligned} \boldsymbol{\tau} &\sim \text{Dirichlet}\left(\frac{\alpha}{K}, ..., \frac{\alpha}{K}\right) \\ \theta_j &\sim H \\ Z_i \mid \boldsymbol{\tau} &\sim \text{Multinomial}\left(\boldsymbol{\tau}, 1\right) \\ X_i \mid (Z_{ij} = 1), \theta_j &\sim f_X\left(X_i \mid \theta_j\right) \end{aligned} \tag{2}$$

$\alpha$ parametrizes the strength of the prior. Higher $\alpha$ generates models that require more trajectories in order for the posterior distribution over $\boldsymbol{\tau}$ to depart from its prior distribution. We emphasize that 2 assumes knowledge of $K$, the number of states.

The denominator in equation 1 is not analytically tractable for mixture models like 2. Solutions to this problem usually fall into one of two categories. The first is to sample from 1 numerically, the approach taken by Markov chain Monte Carlo (MCMC) methods. The second is to construct a tractable approximation to 1, the approach taken by variational Bayes (VB) methods. DPMMs rely on the first approach, while SAs rely on the second. We summarize a simple VB algorithm for finite-state mixtures of Brownian motions in Appendix C, from which a VB algorithm for inference on state arrays can be derived as a special case.

While the full posterior distribution over model parameters is useful in its own right, it is convenient to derive a "working estimate" for the model parameters using some metric on the posterior distribution. The two most common metrics are the the mean of the posterior distribution and the maximum *a posteriori* parameters. Throughout this manuscript, we always use the posterior mean.

#### 1.2 Mixture models for fragmented spaSPT datasets

While attractive, the assumption $Z_i \mid \boldsymbol{\tau} \sim \text{Multinomial}(\boldsymbol{\tau}, 1)$ introduces serious biases when applied to mammalian spaSPT for two reasons:

1. New trajectories not only originate from fluorophores that photoactivate within the focal volume, but also from fluorophores that diffuse into focus from outside. Faster trajectories are more likely to diffuse into focus than slow ones. Consequently, fast states contribute more trajectories than slow states with the same underlying fractional occupancy.

2. The probability to capture a jump requires that both endpoints of the jump lie within focus. The faster the diffusion of a particular state, the more of its jumps will end outside the focus.

How do we address each of these issues? To start with, we can notice that even though the paths of fast particles are fragmented into many short observed trajectories, the probability that a particle in state $j$ is observed inside the focal volume *at any single frame* is $\tau_j$, regardless of its mobility. As a result, the probability to observe a single jump from state $j$ is proportional to $\eta_j \tau_j$, where $\eta_j$ is the probability that the endpoint of the jump lies within focus. $\eta_j$ will generally depend on the state's dynamic parameters $\theta_j$; Appendix B focuses on algorithms to compute $\eta_j$.

In this way, for a dataset with $N$ trajectories, the expected number of jumps contributed from state $j$ is proportional to $\sum_{i=1}^{N} \frac{dL_i}{2} \mathbb{E}[Z_{ij}]$, where $L_i$ is the number of jumps in trajectory $i$, $\mathbb{E}[Z_{ij}] = \Pr(Z_{ij} = 1) \propto \eta_j \tau_j f_X(X_i | \theta_j)$, and $d$ is the number of spatial dimensions. (The factor $d/2$, while unnecessary here as it normalizes out over all the states, is included to highlight similarities with the multivariate normal and gamma likelihoods for trajectories discussed in Appendix A.)

Then we can estimate the true state occupations $\eta_j$ by dividing out the defocalization factor $\eta_j$:

$$\tau_j \propto \frac{\#\text{ of jumps in state } j}{\eta_j} = \frac{\sum_{i=1}^{N} \frac{dL_i}{2} \mathbb{E}[Z_{ij}]}{\eta_j}$$

These relations, which are reminiscent of an expectation-maximization routine, are made concrete in subsequent sections. The important part here is to highlight two deviations of our approach from the classic mixture model represented by 2: (1) we determine state occupations by counting jumps rather than trajectories, and (2) we distinguish between the underlying state occupations $\boldsymbol{\tau}$ and the observed state occupations, which are related through the defocalization factor $\eta_j$.

#### 1.3 Model complexity and localization error

In the context of mammalian spaSPT experiments, there are two additional challenges associated with mixture models like 2. The first is that, in cellular SPT, the number of states $K$ is often unknown. Approaches to address this problem either incorporate $K$ explicitly as a random variable in the model or discriminate between posteriors for different $K$ using criteria like the evidence lower bound (ELBO). The ELBO approach is taken, for example, by the vbSPT method [1].

DPMMs represent an alternative approach that takes the limit $K \to \infty$ and relies on the regularizing property of Bayesian inference to select a sparse subset of states sufficient to explain the observed trajectories. This approach, while attractive in that it imposes no restrictions on the type of mixture models that can be learned, presents computational challenges for complex likelihood functions.

SAs respond to this challenge by recognizing that $K \to \infty$ is unnecessary for most practical purposes, and that choosing a finite model with a large, finite $K$ and a fixed parameter vector $\boldsymbol{\theta}$ retains the desired property of selecting sparse models from more complex alternatives while providing the ability to work with more complex likelihood functions.

The second challenge associated with mixture models for spaSPT is that measurement error often affects each state unequally. Particles that move more quickly have more motion blur, resulting in higher uncertainty in their position ("localization error"). Our preferred approach to address this problem is to incorporate localization error explicitly as a state parameter, then marginalize over it after inference. This procedure, while prohibitively costly for DPMMs, is straightforward for SAs.

#### 2 Dirichlet process mixture models (DPMMs) for trajectory mixtures

DPMMs are the limit case of the finite-state mixture model 2 that results when we take $K \to \infty$. Of course, estimating the parameters for models with large $K$ via maximum likelihood or least-squares approaches leads to hopeless overfitting. For instance, extracting state occupations through jump distribution fitting is only realistic with a handful of states. Fitting a 100-state model to a jump distribution results in many states being adaptations to noise or effectively redundant, splitting a real underlying state into multiple model states with similar parameters. Interpretation of such fits is usually impossible.

Bayesian methods as represented by 2, however, impose an inherent penalty on model complexity via the prior distribution, and this leads them to favor sparse, intelligible models in which only a subset of states are used to "explain" the observed data. Indeed, as $K$ becomes large, the number of states with nonzero occupancy in the posterior for models like 2 becomes *independent* of $K$ and instead grows as $\alpha \log N$, where $\alpha$ is the prior strength and $N$ is the number of observations in the dataset [2].

Inspired by this observation, the DPMM is obtained by letting $K \to \infty$ [6]. In this limit, the complexity of the posterior model - the number of states required to "explain" a set of trajectories - depends solely on the trajectories themselves and not on the researcher's knowledge about the number of states. For many applications in live cells, where the ground truth about the number of states is unknown before doing the experment, this agnostic approach is attractive.

In this section, we first review some properties of DPMMs relevant for their application to trajectories from SPT experiments. Next, we derive a DPMM inference algorithm for mixtures of regular Brownian motions. Finally, we point out some shortcomings of this algorithm, especially its awkward relationship with localization error. These shortcomings are largely addressed by state arrays, which can be considered a DPMM-inspired compromise between the low-$K$ and infinite-$K$ cases.

#### 2.1 Replacing finite state models with Dirichlet processes

Like the regular Dirichlet distribution, Dirichlet processes (DPs) are probability distributions whose draws are themselves probability distributions. However, while draws from a regular Dirichlet distribution are always categorical distributions over a fixed set of $K$ states, each draw from a DP has a distinct, random support [5]. This property is crucial, because it means even though each draw from a DP is a discrete distribution, marginalizing over these draws produces *continuous* distributions over a parameter space of interest - for instance, the diffusion coefficient for regular Brownian motion.

The Dirichlet process is defined in the following way [3]. Suppose $H$ is a measure on a parameter space of interest $\Theta$. The random measure $G$ is distributed according to a Dirichlet process with concentration parameter $\alpha$ and base distribution $H$ (written as $G \sim \text{DP}(\alpha, H)$) if, for any finite partition $A_1, ..., A_K \subseteq \Theta$ of the parameter space,

$$(G(A_1), ..., G(A_K)) \sim \text{Dirichlet}\left(\alpha H(A_1), ..., \alpha H(A_K)\right) \tag{3}$$

Here, $G(A)$ and $H(A)$ are the probability that a draw from $G$ or $H$ falls into the set $A$, respectively. Then the properties of the regular Dirichlet distribution afford us the mean and variance of the draws $G$:

$$\begin{aligned}\mathbb{E}\left[G(A)\right] &= H(A) \\ \text{Var}\left[G(A)\right] &= \frac{H(A)(1 - H(A))}{1 + \alpha}\end{aligned}$$

The base distribution functions like the "mean" of the realizations $G$. That is, the DP generates probability distributions about $H$ similar to how a regular distribution generates random variables around its mean. The variability of these distributions is inversely related to the concentration parameter: as $\alpha \rightarrow \infty$, the draws become close to $H$.

From a Bayesian perspective, $H$ is the prior distribution over the parameters, while selection of the concentration parameter reflects the degree to which we trust incoming data over our naïve expectation. Higher $\alpha$ means the data must work harder to convince us of the existence of new diffusive states.

With these considerations in mind, the Dirichlet process mixture model can be defined [6]

$$\begin{aligned} G &\sim \mathrm{DP}(\alpha, H) \\ \theta_i \mid G &\sim G \\ X_i &\sim f_X(X_i \mid \theta_i) \end{aligned} \tag{4}$$

where $X_i$ is the $i^{\text{th}}$ trajectory in a dataset of $N$ trajectories, and $\theta_i$ are parameters governing its motion according to some PDF $f_X(X_i|\theta_i)$. Although DPMMs may be somewhat daunting at first glance, notice that 4 is arguably simpler than the finite-state mixture model 2. In effect, the DPMM replaces the separate parametrizations of state parameters and occupations in 2 with a single, continuous distribution over all possible state parameters. It corresponds directly to the following generative model: when the microscope observes a trajectory, we draw a random set of parameters $\theta_i$ from an underlying distribution over state parameters, then generate a random trajectory $X_i$ from those parameters. Our goal is to approximate the underlying distribution over $\theta$ by evaluating the conditional distribution $p(\theta|\mathbf{X})$.

#### 2.2 Gibbs samplers for DPMMs

To evaluate the posterior distribution for a DPMM, we take the Gibbs sampling approach introduced by Neal [7]. This involves drawing from the conditional distribution of $\theta_i$ while holding the other $\theta_j$ for which $j \neq i$ constant. (For convenience, we let $\boldsymbol{\theta}_{-i}$ denote the set of all $\theta_j$ for which $j \neq i$, so that we can write this conditional distribution as $p(\theta_i|\mathbf{X}, \boldsymbol{\theta}_{-i})$.) Cycling through all $\theta_i$ this way, the sequence of $\theta$ thus produced constitutes a sample from the posterior distribution $p\left(\theta|\mathbf{X}\right)$ [8].

Building on observations from Blackwell & MacQueen [4], Neal showed that this conditional distribution follows the relation

$$p\left(\theta_i|\mathbf{X}, \boldsymbol{\theta}_{-i}\right) \propto p\left(X_i|\theta_i\right) \left[ \frac{\alpha H(\theta_i) + \sum\limits_{j \neq i} \delta_{\theta_j}(\theta_i)}{\alpha + N - 1} \right] \tag{5}$$

Here, $H(\theta_i)$ is the probability density at $\theta_i$ under the prior, while each $\delta_{\theta_i}$ is an identity probability measure: $\delta_{\theta_i}(A) = 1$ if $\theta_i \in A$ and 0 otherwise. The term $p(X_i|\theta_i)$ is just the PDF for our diffusion model, to be selected shortly. The term in brackets on the right side is analogous to the role that the state occupations $\boldsymbol{\tau}$ play in the FSMM (Appendix C). Altogether, sampling from $p(\theta_i \mid \mathbf{X}, \boldsymbol{\theta}_{-i})$ can be accomplished in the following way:

> Draw a random number $u \sim \text{Uniform}(0, 1)$. Depending on the result, do one of the following:

- If $u \leq \frac{\alpha}{\alpha+N-1}$, draw $\theta_i$ with probability proportional to $p(X_i|\theta_i)H(\theta_i)$.
- If $u > \frac{\alpha}{\alpha+N-1}$, set $\theta_i$ equal to one of the other values in $\boldsymbol{\theta}_{-i}$ with probability proportional to the number of times that value occurs in $\boldsymbol{\theta}_{-i}$, multiplied by $p(X_i|\theta_i)$.

Notice that the vector $\boldsymbol{\theta} = (\theta_1, ..., \theta_N)$ changes at each iteration as it is gradually "shuffled" by these state reassignments. A complete cycle through all $N$ trajectories produces some vector $\boldsymbol{\theta}^{(t)}$ ($t$ standing for the $t^{\text{th}}$ iteration) that is a finite sample from the posterior distribution $p(\theta|\mathbf{X})$. $\boldsymbol{\theta}^{(t)}$ is then used as the starting point for the $(t+1)^{\text{th}}$ iteration, and the process continues.

Although this is a viable Gibbs sampler for DPMMs, it is inefficient, exploring the posterior slowly. A large number of trajectories assigned to some favorable $\theta$ can function as a "sink" from which it can take many iterations to escape. In response, Neal added an additional Metropolis-Hastings step to nudge the parameters in $\boldsymbol{\theta}$ by a little after each round of shuffling (Algorithm 8 in [7]). The stationary distribution of this modified Gibbs sampler still corresponds to the posterior distribution $p(\theta|\mathbf{X})$, but it generates representative samples with fewer iterations than the naïve sampler. In what follows, we follow this scheme with Gaussian nudges.

#### 2.3 Gibbs sampler for RBME

We apply Neal's Gibbs sampler to the case of regular Brownian motion with localization error (RBME). As discussed in detail in Appendix A, RBME is a stochastic model in which the apparent "jump" of a particle from time $t$ to time $t + \Delta t$ has contributions from both the true underlying motion of the particle and *localization error*, the imprecision associated with measuring a particle's position. Specifically, the apparent 1D jumps of an RBME have mean zero and variance $2(D\Delta t + \sigma^2_{\text{loc}})$, where $D$ is the diffusion coefficient and $\sigma^2_{\text{loc}}$ is the variance of the localization error.

Localization error not only affects the apparent jump magnitude, but also induces nonzero covariance between subsequent jumps in a trajectory. Neglecting these covariances leads to the *gamma approximation* for the RBME likelihood, which is

$$f_X(X_i|\phi_j) = \frac{S_i^{\frac{dL_i}{2}-1} e^{-S_i \phi_j^{-1}}}{\Gamma\left(\frac{dL_i}{2}\right) \phi_j^{\frac{dL_i}{2}}}$$

where $S_i$ is the sum of squared jumps for trajectory $i$, $L_i$ is the number of jumps in the trajectory, $d$ is the number of spatial dimensions, and $\phi_j = 4(D_j \Delta t + \sigma_j^2)$ parametrizes the variance of the jumps for state $j$. The simple form of the gamma likelihood means that we can succinctly represent the $i^{\text{th}}$ trajectory $X_i$ as a tuple of two numbers: $X_i = (S_i, L_i)$. Notice that we cannot distinguish the contributions of $D_j$ and $\sigma_j^2$ to $\phi_j$ without measuring $\sigma_j^2$ by some other method, such as averaging the negative sequential jump covariance across all trajectories in a dataset - this is the price we pay for a tractable likelihood function.

As discussed in section 1.2, the equation above neglects the effect of defocalization. Specifically, the jumps of each state $j$ have an associated probability $\eta_j$ to be observed given the microscope's focal depth. The true occupation $\tau_j$ of state $j$ is related to its *apparent* occupation $\tau_j^{(\text{obs})}$ as $\tau_j \propto \tau_j^{(\text{obs})} \eta_j^{-1}$. (Algorithms to calculate $\eta_j$ are presented in Appendix B.) So when shuffling trajectories according to 5, we weight the probability to assign a trajectory to each state by $\eta_j^{-1}$.

The corresponding log likelihood, neglecting terms that are constant across different states, is

$$\log f_X(X_i|\phi_j) \propto -S_i \phi_j^{-1} - \frac{dL_i}{2} \log \phi_j$$

To make it easier to examine a broader range of diffusion coefficients, it is often convenient to deal with the log jump variance $\omega_j = \log \phi_j$ rather than $\phi_j$ itself. Then the log likelihood becomes

$$\log f_X(X_i|\omega_j) \propto -S_i e^{-\omega_j} - \frac{dL_i}{2} \omega_j \tag{6}$$

For the prior over $\omega$, we choose a uniform distribution over some range $[\omega_{\text{min}}, \omega_{\text{max}}]$, which we set so that $D_{\text{min}} = 0.01$ $\mu\text{m}^2$ $\text{s}^{-1}$ and $D_{\text{max}} = 100$ $\mu\text{m}^2$ $\text{s}^{-1}$. When assigning trajectory $i$ a parameter randomly sampled from the prior, we follow Neal [7] and first choose $m_0$ random values chosen with uniform probability on the interval $[\omega_{\text{min}}, \omega_{\text{max}}]$. Among these, choose a particular value $\omega'$ with probability proportional to the likelihood of $\omega'$ given trajectory $i$.

For the Metropolis-Hastings updates to the state parameters, we propose updates to each parameter using a mean-zero Gaussian. Specifically, we draw

$$\omega_j' \sim g(\omega|\omega_j^{(t-1)}) = \frac{e^{-(\omega - \omega_j^{(t-1)})^2 / 2\nu^2}}{\sqrt{2\pi\nu^2}}$$

where $\omega_j^{(t-1)}$ is the previous log jump variance for state $j$ and $\nu^2$ is the variance of the proposal distribution.

The final Gibbs sampler for spaSPT DPMMs is the following:

1. Draw a random sample $\boldsymbol{\omega}^{(0)} = (\omega_1, ..., \omega_{m_0})$ from a uniform distribution on the interval $[\omega_{\text{min}}, \omega_{\text{max}}]$. Each element of the vector $\boldsymbol{\omega}^{(0)}$ represents a candidate "state".

2. Assign each trajectory $i$ to the $j^{\text{th}}$ element in $\boldsymbol{\omega}^{(0)}$ with log probability proportional to $-S_i e^{-\omega_j} - \frac{dL_i}{2} \omega_j - \log \eta_j$. Let this assignment be $Z_i^{(0)}$.

3. For each iteration $t = 1, 2, ...$

(a) For each trajectory $i = 1, 2, ...$, either set $Z_i^{(t)}$ to a state in the current set $\boldsymbol{\omega}^{(t-1)}$ with probability $(N-1)/(\alpha+N-1)$, or create a new state with probability $\alpha/(\alpha+N-1)$.

   i. If setting to an existing state, choose state $j$ with log probability proportional to $\log n_j - \log \eta_j + \log f_X(X_i|\omega_j)$, where $n_j$ is the number of jumps already assigned to state $j$ and $\eta_j$ is the probability that a jump from state $j$ is observed after accounting for defocalization.

   ii. If creating a new state, pick $m_0$ values of $\omega$ from the interval $[\omega_{\min}, \omega_{\max}]$. Among these, accept a particular value $\omega'$ with log probability proportional to $\log f_X(X_i|\omega')$. Add this new state $\omega'$ to the set of current states $\boldsymbol{\omega}^{(t-1)}$.

(b) For each state $j$, if there are no trajectories currently assigned to it, remove it from consideration. Otherwise add it to $\boldsymbol{\omega}^{(t)}$, the next set of states, and update it according to a Metropolis-Hastings step as follows:

   i. Propose a new $\omega' \sim g(\omega' \mid \omega_j^{(t-1)})$ and evaluate the likelihood ratio

$$r = \frac{\prod_{i=1}^{N} \mathbb{I}_{Z_i=j} f_X(X_i|\omega')}{\prod_{i=1}^{N} \mathbb{I}_{Z_i=j} f_X(X_i|\omega_j^{(t-1)})} \frac{\Phi\left(\frac{\omega_{\max}-\omega_j^{(t-1)}}{\nu}\right) - \Phi\left(\frac{\omega_{\min}-\omega_j^{(t-1)}}{\nu}\right)}{\Phi\left(\frac{\omega_{\max}-\omega'}{\nu}\right) - \Phi\left(\frac{\omega_{\min}-\omega'}{\nu}\right)}$$

   ii. Draw $u \sim \text{Uniform}(0,1)$. If $r > u$, set $\omega_j^{(t)} = \omega_j^{(t-1)}$. Otherwise set $\omega_j^{(t)} = \omega'$.

4. Return the set of all samples $\boldsymbol{\omega}^{(t)}$ and $\mathbf{n}^{(t)}$, where $n_j^{(t)}$ is the total number of jumps assigned to state $j$ at iteration $t$.

In this algorithm, $\Phi(x)$ is the unit Gaussian CDF; its contribution to $r$ is required to make an unbiased proposal distribution over the range $[\omega_{\min}, \omega_{\max}]$, necessary for the ergodicity of the Gibbs sampler.

As mentioned previously, we can make an estimate of the posterior distribution by taking a histogram of the samples $\boldsymbol{\omega}^{(t)}$ weighted by their occupations $\mathbf{n}^{(t)}$.

While the gamma approximation is what makes DPMMs computationally tractable, it also means that in order to disambiguate the contributions of diffusion and localization error to $\omega$ we need to measure localization error by a different method. This is particularly relevant for calculating the defocalization factor $\eta_j$, which relies on knowledge of $D_j$ independent of $\sigma^2$ (Appendix B). In this manuscript, we always use the mean negative covariance between subsequent jumps to estimate localization error prior to launching a DPMM, so that $\eta_j$ can be estimated from $D_j$. However, this means that our DPMM is only as good as our estimate of $\sigma^2$ - and as demonstrated in Fig. 3 and S3, our estimate of $\sigma^2$ can be quite noisy with small numbers of trajectories, and starts to fail completely when localization error varies a lot between states. In contrast, state arrays provide a more elegant approach to disambiguate the effects of localization error and diffusion.

### 3 State arrays

A disadvantage of the DPMM is its high computational cost. In order to execute the algorithm above, we need to evaluate the likelihood $f_X(X_i|\omega_j)$ for every trajectory $i$ and every active state $j$ at each iteration $t$. This is feasible for the log gamma likelihood 6 because of its simplicity, but becomes costly for more complex likelihood functions. For instance, if we want to remove the assumption that localization error is the same for all states - and this is highly desirable - then the DPMM must infer posterior occupations over the joint parameter space of diffusion coefficient and localization error, an impractical task for most computers.

We can replace the DPMM with a different model that retains many of the DPMM's attractive properties - such as its ability to model complex mixtures - while mitigating the issue of expensive likelihood functions. To do this, we return to the finite-state model 2. Instead of letting the number of states $K \to \infty$, we instead fix $K$ at a large but finite value. For each state $j = 1, ..., K$, choose a fixed set of state parameters $\theta_j$. For instance, we may choose the set of all $\theta_j$ as an evenly spaced grid spanning the target region of parameter space.

As a result, model 2 simplifies to

$$\begin{aligned} \boldsymbol{\tau} &\sim \text{Dirichlet}\left(\frac{\alpha}{K}, ..., \frac{\alpha}{K}\right) \\ Z_i \mid \boldsymbol{\tau} &\sim \text{Multinomial}\left(\boldsymbol{\tau}, 1\right) \\ X_i \mid (Z_{ij} = 1) &\sim f_X(x \mid \theta_j) \end{aligned} \tag{7}$$

Notice that since each $\theta_j$ is invariant, $f_X(x \mid \theta_j)$ is also invariant and only needs to be evaluated once at the beginning of inference.

#### 3.1 Posterior evaluation

In the context of finite-state mixture models like 2, the assumption of static $\theta_j$ can be implemented by replacing the prior over $\theta_j$ with a Dirac delta function. As a result, we can adapt the VB inference scheme for FSMMs outlined in Appendix C as follows.

The goal of inference is to evaluate the posterior $p(\mathbf{Z}, \boldsymbol{\tau} \mid \mathbf{X})$. We approximate the posterior with a function $q(\mathbf{Z}, \boldsymbol{\tau})$ such that

$$q(\mathbf{Z}, \boldsymbol{\tau}) = q(\mathbf{Z})q(\boldsymbol{\tau})$$
$$q = \underset{q'}{\operatorname{argmax}}\, L[q']$$

where $L[q]$ is the evidence lower bound (ELBO):

$$L[q] = -\int q(\mathbf{Z}, \boldsymbol{\tau}) \log\left[\frac{q(\mathbf{Z}, \boldsymbol{\tau})}{p(\mathbf{X}, \mathbf{Z}, \boldsymbol{\tau})}\right] \mathrm{d}\mathbf{Z}\, \mathrm{d}\boldsymbol{\tau}$$

A justification for this scheme is briefly reviewed in Appendix C and can be found in any text on variational Bayesian methods. Maximization of $L[q]$ can be accomplished by alternately evaluating

$$\log q(\mathbf{Z}) = \mathbb{E}_{\boldsymbol{\tau}\sim q(\boldsymbol{\tau})}\left[\log p\left(\mathbf{X}, \mathbf{Z}, \boldsymbol{\tau}\right)\right] + \text{constant}$$
$$\log q(\boldsymbol{\tau}) = \mathbb{E}_{\mathbf{Z}\sim q(\mathbf{Z})}\left[\log p\left(\mathbf{X}, \mathbf{Z}, \boldsymbol{\tau}\right)\right] + \text{constant}$$

The constants are chosen so that each factor of $q$ is normalized.

First, we solve $q(\boldsymbol{\tau})$. Note that we can factor $p(\mathbf{X}, \mathbf{Z}, \boldsymbol{\tau})$ as $p(\mathbf{X}|\mathbf{Z}, \boldsymbol{\tau})p(\mathbf{Z}|\boldsymbol{\tau})p(\boldsymbol{\tau})$. For $p(\boldsymbol{\tau})$, we choose the Dirichlet prior $p(\boldsymbol{\tau}) = \text{Dirichlet}\left(\frac{\alpha}{K}, ..., \frac{\alpha}{K}\right)$. Then

$$\log q(\boldsymbol{\tau}) = \sum_{j=1}^{K}\left(\frac{\alpha}{K} - 1 + \sum_{i=1}^{N}\frac{dL_i}{2}\mathbb{E}_{\mathbf{Z}\sim q(\mathbf{Z})}\left[Z_{ij}\right]\right)\log \tau_j + \text{constant}$$
$$q(\boldsymbol{\tau}) = \text{Dirichlet}\left(\frac{\alpha}{K} + \sum_{i=1}^{N}\frac{dL_i}{2}\mathbb{E}_{\mathbf{Z}\sim q(\mathbf{Z})}\left[Z_{i,1}\right], ..., \frac{\alpha}{K} + \sum_{i=1}^{N}\frac{dL_i}{2}\mathbb{E}_{\mathbf{Z}\sim q(\mathbf{Z})}\left[Z_{i,K}\right]\right)$$

Next, we solve for $q(\mathbf{Z})$. Using the same factorization, we have

$$\log q(\mathbf{Z}) = \sum_{i=1}^{N}\sum_{j=1}^{K} Z_{ij}\left(\mathbb{E}_{\boldsymbol{\tau}\sim q(\boldsymbol{\tau})}\left[\log \tau_j\right] + \log f_X(X_i|\theta_j)\right) + \text{constant}$$

Given that $q(\boldsymbol{\tau})$ is a Dirichlet distribution,

$$\mathbb{E}_{\boldsymbol{\tau}\sim q(\boldsymbol{\tau})}\left[\log \tau_j\right] = \psi(n_j) - \psi\left(\sum_{k=1}^{K} n_k\right)$$

where

$$n_j = \frac{\alpha}{K} + \sum_{i=1}^{N}\frac{dL_i}{2}\mathbb{E}_{\mathbf{Z}\sim q(\mathbf{Z})}\left[Z_{ij}\right]$$

and $\psi(x) = \mathrm{d}\Gamma(x)/\mathrm{d}x$ is the digamma function. As a result,

$$q(\mathbf{Z}) = \prod_{i=1}^{N}\prod_{j=1}^{K} r_{ij}^{Z_{ij}}$$
$$r_{ij} = \frac{\rho_{ij}}{\sum\limits_{k=1}^{K}\rho_{ik}}$$
$$\rho_{ij} = f_X(X_i|\theta_j)e^{\psi(n_j)}$$

The final posterior approximation is

$$\begin{aligned}
q(\boldsymbol{\tau}) &= \text{Dirichlet}\left(\frac{\alpha}{K} + \sum_{i=1}^{N}\frac{dL_i}{2}\mathbb{E}_{\mathbf{Z}\sim q(\mathbf{Z})}\left[Z_{i,1}\right], ..., \frac{\alpha}{K} + \sum_{i=1}^{N}\frac{dL_i}{2}\mathbb{E}_{\mathbf{Z}\sim q(\mathbf{Z})}\left[Z_{i,K}\right]\right)\\
q(\mathbf{Z}) &= \prod_{i=1}^{N}\prod_{j=1}^{K} r_{ij}^{Z_{ij}}\\
r_{ij} &= \frac{\rho_{ij}}{\sum\limits_{k=1}^{K}\rho_{ik}}\\
\rho_{ij} &= f_X(X_i|\theta_j)e^{\psi(n_j)}
\end{aligned} \tag{8}$$

For inference, we alternate between evaluating $q(\boldsymbol{\tau})$ and $q(\mathbf{Z})$ according to the following scheme:

1. Initialize:
   (a) Evaluate $f_X(X_i|\theta_j)$ for each trajectory $i$ and each state $j$. Represent these likelihoods as an $N$-by-$K$ matrix $\mathbf{A}$.
   (b) Set $r_{ij}^{(0)} = A_{ij}/\sum_{k=1}^{K} A_{ik}$.
2. For each iteration $t = 1, 2, ...$
   (a) For each state $j$, evaluate $n_j^{(t)} = \frac{\alpha}{K} + \sum_{i=1}^{N}\frac{dL_i}{2}r_{ij}^{(t-1)}$.
   (b) Evaluate the matrix $\mathbf{r}^{(t)}$ such that

$$r_{ij}^{(t)} = \frac{A_{ij}e^{\psi(n_j^{(t)})}}{\sum\limits_{k=1}^{K} A_{ik}e^{\psi(n_k^{(t)})}}$$

3. After convergence, the posterior distribution can be summarized with the posterior mean:

$$\mathbb{E}\left[\tau_j\right] = \frac{n_j}{\sum\limits_{k=1}^{K} n_k}$$

$$\mathbb{E}\left[Z_{ij}\right] = r_{ij}$$

As discussed in section 1.2, the probability $\eta_j$ to observe a jump from state $j$ is dependent on the state parameters $\theta_j$ due to the influence of defocalization. This leads to systematic overestimates for the occupations of slow states. While it is possible to incorporate $\eta_j$ directly into the inference routine above, we find that it makes little difference if we incorporate it as a final polish step for the posterior mean, so that

$$\mathbb{E}_{\text{defoc}}\left[\tau_j\right] = \frac{n_j / n_j}{\sum\limits_{k=1}^{K} n_k / \eta_k}$$

where, as before, $\mathbf{n} = (n_1, ..., n_k)$ is the parameter for the Dirichlet posterior $q(\boldsymbol{\tau})$.

#### 3.2 Special case: regular Brownian motion with localization error

So far in our discussion of state arrays, we have not specified the form of the likelihood function $f_X(x|\theta)$. State arrays are useful in that, because they only require evaluation of the likelihood function once at the beginning of the algorithm, they are practical for use with likelihood functions that are too costly for DPMMs.

Consider a state array for regular Brownian motion with localization error (Appendix A), a dynamic model in which each state is characterized by two parameters: a diffusion coefficient $D$ and a localization error $\sigma^2_{\text{loc}}$. If trajectory $i$ has $L_i$ jumps, and we use $\mathbf{x}_i, \mathbf{y}_i \in \mathbb{R}^{L_i}$ to represent the vectors of 1D jumps along the $x$ and $y$ axes, respectively, then the RBME likelihood is given by eq. 10. There are two parameters for each state: a diffusion coefficient $D$ and a localization error $\sigma^2_{\text{loc}}$.

In this case, we can choose the states to span a grid of values of $D$ and $\sigma^2_{\text{loc}}$. For example, let $\boldsymbol{\theta}_D = (D_1, ..., D_n)$ be a vector of diffusion coefficients and $\boldsymbol{\theta}_{\sigma^2_{\text{loc}}} = \left(\sigma^2_{\text{loc},1}, ..., \sigma^2_{\text{loc},m}\right)$ be a vector of localization errors. Then we choose the state parameters $\boldsymbol{\theta}$ as the Cartesian product of $\boldsymbol{\theta}_D$ and $\boldsymbol{\theta}_{\sigma^2_{\text{loc}}}$. In the main text, we often use a grid of $n = 201$ diffusion coefficients that are log-spaced between 0.01 and 100 $\mu\text{m}^2\ \text{s}^{-1}$ and $m = 61$ localization errors with root variance between 0 and 100 $\mu$m, for a total of $K = nm = 12060$ states.

After obtaining the posterior distribution $\boldsymbol{\tau}$ over the state occupations, we can then marginalize on localization error by summing the elements of $\boldsymbol{\tau}$ that correspond to each unique diffusion coefficient.

### A Appendix: Regular Brownian motion with localization error

This appendix reviews regular Brownian with localization error (RBME), the underlying stochastic model for the models considered in this manuscript.

#### A.1 Definition

We use *regular Brownian motion (RBM) in one dimension* to refer to a stochastic process $B_t$ such that

$$\begin{aligned} B_0 &= 0 \\ B_{t+\Delta t} - B_t &\sim \mathcal{N}(0, 2D\Delta t) \\ \mathrm{Cov}\left(B_{t+\Delta t} - B_t, B_{s+\Delta s} - B_s\right) &= 0 \end{aligned}$$

where $[t, t+\Delta t]$ and $[s, s+\Delta s]$ are non-overlapping time intervals.

Equivalently, we can define $B_t$ as a Gaussian process with the covariance function $\mathrm{Cov}(B_t, B_s) = 2D \cdot \min(t, s)$.

RBM in multiple spatial dimensions is generated by combining independent RBMs along each spatial dimension. In a single spatial dimension with zero localization error, RBM coincides with the Wiener process.

RBM should be distinguished from phenomenological Brownian motion, the erratic movement of microscopic objects in fluids without net advection. RBM, first defined in a physical context by Einstein [9], serves as a model for phenomenological Brownian motion in low Reynolds number fluids under conditions when the forces on a microscopic object are numerous and uncorrelated enough to apply the central limit theorem to the resulting net force vector.

#### A.2 Localization error

Even when RBM is a good model for the underlying motion, we never measure $B_t$ exactly due to the ubiquity of *localization error*, the imprecision associating with measuring a particle's position. This is particular relevant for spaSPT experiments, in which the localization error can be similar in magnitude to the size of the jumps themselves.

To model localization error, let $E_t$ be a Gaussian white noise process, so that

$$\begin{aligned}\mathbb{E}\left[E_t\right] &= 0 \\ \mathrm{Cov}(E_t, E_s) &= \sigma_{\mathrm{loc}}^2 \delta(t-s)\end{aligned}$$

where $\delta(t-s)$ is the Dirac delta function. Then we define regular Brownian motion with localization error (RBME) in one dimension as

$$\overline{B}_t = B_t + E_t$$

$\overline{B}_t$ is another Gaussian process such that

$$\begin{aligned}\mathbb{E}\left[\overline{B}_t\right] &= 0 \\ \mathrm{Cov}\left(\overline{B}_t, \overline{B}_s\right) &= 2D \cdot \min(t,s) + \sigma_{\mathrm{loc}}^2 \delta(t-s)\end{aligned}$$

#### A.3 Increments

In spaSPT, we measure the coordinates of a particle at regular frame intervals of length $\Delta t$. For this case, we can define the jump process $\tilde{B}_n = \overline{B}_{n\Delta t} - \overline{B}_{(n-1)\Delta t}$. This is another mean-zero Gaussian process. Using linearity of expectation, we can see that

$$\begin{aligned}\mathrm{Cov}\left(\tilde{B}_n, \tilde{B}_m\right) &= \mathbb{E}\left[\tilde{B}_n \tilde{B}_m\right] \\ &= \mathbb{E}\left[\left(\overline{B}_{n\Delta t} - \overline{B}_{(n-1)\Delta t}\right)\left(\overline{B}_{m\Delta t} - \overline{B}_{(m-1)\Delta t}\right)\right] \\ &= \mathrm{Cov}\left(\overline{B}_{n\Delta t}, \overline{B}_{m\Delta t}\right) - \mathrm{Cov}\left(\overline{B}_{n\Delta t}, \overline{B}_{(m-1)\Delta t}\right) \\ &\quad - \mathrm{Cov}\left(\overline{B}_{(n-1)\Delta t}, \overline{B}_{(m-1)\Delta t}\right) + \mathrm{Cov}\left(\overline{B}_{(n-1)\Delta t}, \overline{B}_{(m-1)\Delta t}\right) \\ &= \begin{cases} 2\left(D\Delta t + \sigma_{\mathrm{loc}}^2\right) & \text{if} \quad m = n \\ -\sigma_{\mathrm{loc}}^2 & \text{if} \quad |m-n| = 1 \\ 0 & \text{otherwise} \end{cases}\end{aligned} \tag{9}$$

Importantly, the covariance is nonzero for sequential jumps in a trajectory (jumps that share a common point).

Using 9, we can write out the PDF for RBME explicitly. Suppose the $i^{\mathrm{th}}$ trajectory in a dataset has $L_i$ jumps. One way to represent this trajectory is a tuple $X_i = (\mathbf{x}_i, \mathbf{y}_i)$ where $\mathbf{x}_i, \mathbf{y}_i \in \mathbb{R}^{L_i}$ are the vectors of its 1D jumps along the $x$ and $y$ axes respectively. Then the joint PDF for $\mathbf{x}_i$ and $\mathbf{y}_i$ is

$$f_{X_i}(\mathbf{x}_i, \mathbf{y}_i \mid D, \sigma_{\mathrm{loc}}^2) = \frac{\exp\left(-\frac{1}{2}\left[\mathbf{x}_i^T \Sigma^{-1} \mathbf{x}_i + \mathbf{y}_i^T \Sigma^{-1} \mathbf{y}_i\right]\right)}{2\pi \det(\Sigma)} \tag{10}$$

where $\Sigma_{ij} = \mathrm{Cov}\left(\tilde{B}_i, \tilde{B}_j\right)$ as given by 9.

##### A.4 Gamma approximation

If the off-diagonal terms in the covariance 9 can be neglected, then the process can be characterized by a single parameter $\phi_j = 4(D\Delta t + \sigma_{\text{loc}}^2)$. In this case, the multivariate normal PDF 10 is equivalent to the gamma density

$$f_X(X_i|\phi_j) = \begin{cases} \frac{S_i^{\frac{dL_i}{2}-1} e^{-S_i \phi_j^{-1}}}{\Gamma\left(\frac{dL_i}{2}\right) \phi_j^{\frac{dL_i}{2}}} & \text{if } \phi_j \geq 0 \\ 0 & \text{otherwise} \end{cases} \tag{11}$$

where $S_i = \sum_{n=1}^{L_i} (\mathbf{x}_i)_n^2 + (\mathbf{y}_i)_n^2$ is the sum of squared jumps for the trajectory. In this case, the trajectory $X_i$ can be represented succinctly as the tuple $(S_i, L_i)$.

### B Appendix: Defocalization

spaSPT experiments are typically acquired at frame intervals of several milliseconds. $z$-stacks are unfeasible at these speeds. Combined with the short focal depths induced by the high numerical aperture objectives used for tracking experiments, observation of fluorescent particles is typically confined to a thin observation slice ("focal volume") less than a micron in depth [14]. Tracking in multiple focal planes simultaneously is a promising approach to mitigate this problem [15], but such approaches are not yet widely accessible.

The rate of defocalization was considerd as a way to measure the diffusion coefficient by Kues & Kubitscheck [16] and was incorporated as a correction term into jump histogram modeling frameworks by Mazza [12] and Hansen & Woringer [13]. Whether it is considered bug or feature, it plays a fundamental role in coupling the mobility characteristics of each state with the state's apparent occupation, and it cannot be neglected for spaSPT experiments with short focal depth.

[12] and [13] evaluated the defocalization probability by using the solution to the diffusion equation within a slab with absorbing boundaries [17]. Because the boundaries for the focal volume are not actually absorbing, both sets of authors then applied a correction term derived from Monte Carlo simulations of regular Brownian motion. Here, we provide a simpler alternative that is not based on Monte Carlo simulations. Some of the attractive characteristics of this alternative are that it can also incorporate non-uniform probabilities of detection in the axial direction, works for any number of gaps allowed during tracking, and extends to a broader class of diffusion processes than regular Brownian motion.

In this section, we first review the approach for gapless tracking, then extend it to gapped tracking.

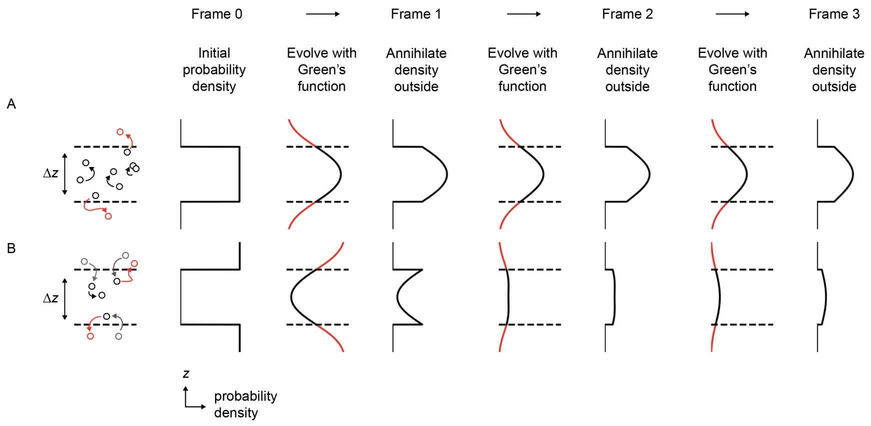

**Figure 1: Approach to calculate defocalized fraction for a Markov diffusion process.** At each iteration, the axial probability density for the particle is convolved with its Green's function, then the "unobserved" probability density is set to zero. (A) and (B) depict two possible initial profiles for the axial density, for a particle that either starts outside or inside the focal volume.

#### B.1 Gapless tracking

In spaSPT we observe the particle's position at a discrete set of timepoints separated by some frame interval $\Delta t$. If we use a gapless tracking algorithm (an algorithm that returns trajectories with no gap frames), then if a particle is not detected a single frame interval, it is lost for all subsequent frame intervals.

Let $f(z, t = 0)$ be the starting profile of particles in the axial direction of the microscope, let $g(z, \Delta t)$ be the Green's function for the diffusion process at this frame interval, and let $\phi(k, \Delta t) = \mathcal{F}[g]$ be the Fourier transform of $g$. For instance, the Green's function for regular Brownian motion is $\phi(k, \Delta t) = e^{-Dk^2 \Delta t}$. More generally, the Green's function for Levy flights with stability parameter $\alpha$ is $\phi(k, \Delta t) = e^{-D|k|^{\alpha} \Delta t}$.

Then the axial probability density for the particle after one frame interval can be obtained by convolving its initial profile with the Green's function for the diffusion model:

$$f(z, \Delta t) = \mathcal{F}^{-1}\left[\phi(k, \Delta t)\mathcal{F}\left[f(z, 0)\right]\right]$$

To evaluate the fraction of particles lost due to defocalization, we can multiply this density with an appropriate transmission function. For example, if our focal volume is a slab with depth $\Delta z$, infinite XY extent, and perfect recall at any point inside the slab (that is, all particles inside the slab are detected and no particles outside are detected), then our transmission function $T$ is

$$T(z) = \begin{cases} 1 & \text{if } z \in \left[-\frac{\Delta z}{2}, \frac{\Delta z}{2}\right] \\ 0 & \text{otherwise} \end{cases}$$

(This is the transmission function considered by [12] and [13].) Then the fraction of defocalized particles is

$$\text{fraction defoclized after one frame interval} = \int_{-\infty}^{+\infty} T(z) f(z, \Delta t) \mathrm{d}z$$

In the case of regular Brownian motion, this also has a simpler form [16] [17]:

$$\frac{1}{2\Delta z} \int_{-\Delta z/2}^{+\Delta z/2} \left[ \operatorname{erfc}\left( \frac{\frac{\Delta z}{2} - z_0}{\sqrt{4D\Delta t}} \right) + \operatorname{erfc}\left( \frac{\frac{\Delta z}{2} + z_0}{\sqrt{4D\Delta t}} \right) \right] \mathrm{d}z$$

However, this form cannot be used to evaluate the probability to defocalize after *two* frame intervals because this equation "counts" particles that are outside the focal volume at time $\Delta t$ and return to the focal volume at time $2\Delta t$. In gapless tracking, these trajectories are dropped and do not contribute to the fraction of particles remaining in the focal volume.

To account for this reentry effect, we can remove the fraction of particles that are dropped by multiplying with the transmission function, then reconvolving the resulting density with the Green's function to propagate it to the next frame. This can be done any number of times, so that the axial probability density for the particles after $m$ frame intervals is

$$f(z, m \text{ frame intervals}) = \text{Propagate}^{(m)} \left[ f(z', 0) \right](z) \tag{12}$$

where we use $\text{Propagate}^{(m)}$ to denote $m$ sequential applications of the operator

$$\text{Propagate}[f](z) = T(z) \mathcal{F}^{-1} \left[ \phi(k, \Delta t) \mathcal{F}[f] \right]$$

This scheme is illustrated in Fig. 1. The procedure is straightforward to implement on a grid approximation with any scientific computing library. The fraction of particles remaining after $m$ frame intervals is just the sum of this density.

#### B.2 Gapped tracking

When using tracking algorithms that allow gap frames, particles can escape from observations for one or more frames before returning. A recursive scheme to account for these effects is depicted in Fig. 2. At each frame interval, the probability density is split into observed and unobserved components, and these

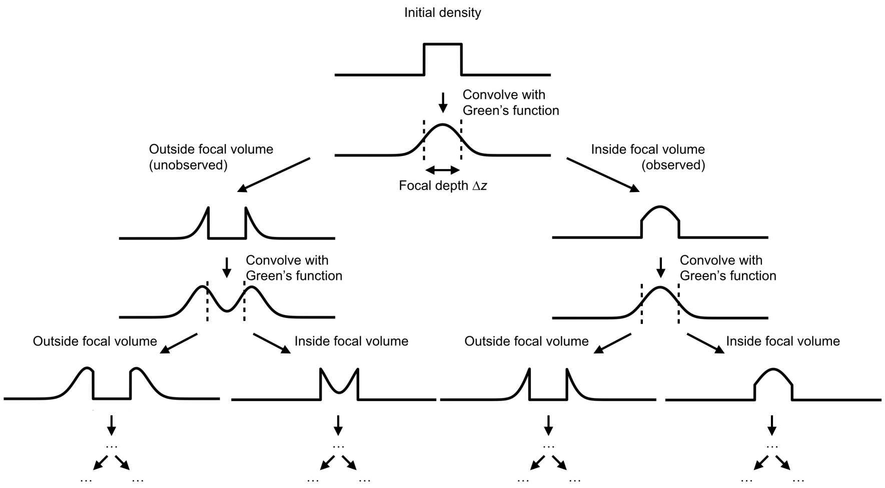

**Figure 2: Recursive approach to calculate defocalized fraction of a Markov process tracked with gaps.** After propagation, the axial probability density for a particle is split into observed and unobserved parts, which are then propagated separately to account for reentry. While here we show a slab observation geometry (uniform detection profile in $z$) for convenience, the algorithm can be adapted to any transmission function.

are propagated separately. While Fig. 2 depicts the transmission function as a slab, we can use this scheme with any axial detection profile by splitting the probability density into parts $Tf$ and $(1-T)f$.

This recursive algorithm can be made iterative if the tracking algorithm has some maximum number of tolerated gaps $M_{\text{gaps}}$. In that case, we can aggregate all of the observed density at each frame interval, while maintaining a buffer of $M_{\text{gaps}}$ distinct unobserved densities. In detail, this algorithm is:

1. Start with an initial profile $f(z, 0)$.
2. Instantiate a buffer for each gap frame: $f_0(z)$, $f_1(z)$, ..., $f_{M_{\text{gaps}}}(z)$. $f_0(z)$ is the observed density, and each subsequent $f_m(z)$ is the density that has been unobserved for $m$ frames. Set $f_0(z) = f(z, 0)$.
3. Instantiate an auxiliary buffer $f_{0,\text{agg}}(z)$.
4. For each frame interval $n = 1, 2, ...$
   (a) Set $f_{0,\text{agg}}(z) = 0$.
   (b) For $m = M_{\text{gaps}}, M_{\text{gaps}} - 1, ..., 0$:
      i. Evolve the probability density $f_m(z) = \text{Propagate}\left[f_m(z)\right]$.

ii. Split the probability density into observed and unobserved parts:

$$f_{m,\text{obs}}(z) = T(z) f_m(z)$$
$$f_{m,\text{unobs}}(z) = (1 - T(z)) f_m(z)$$

iii. Add $f_{m,\text{obs}}(z)$ to $f_{0,\text{agg}}(z)$.

iv. If $m < M_{\text{gaps}}$, set $f_{m+1}(z) = f_{m,\text{unobs}}(z)$.

(c) Set $f_0(z) = f_{0,\text{agg}}(z)$.

The sum of $f_0(z)$ at each frame interval is the fraction of trajectories that are inside the focal volume.

### C Appendix: Variational Bayesian inference for finite mixtures of RBMEs

Here, we outline a variational Bayes (VB) algorithm for finite-state mixture models (FSMMs) like 2. This serves to motivate the use of VB inference for the trajectory state occupation problem and as a precursor to the VB algorithm for state arrays, which is a special case of the algorithm here. The development is closely related to classic expositions of the VB algorithms for mixture models; e.g. in chapter 10 in Bishop's book [10].

#### C.1 Statement of problem

As before, let $\mathbf{X}$ represent a dataset of $N$ trajectories. Each trajectory is associated with some state $j$, and each state $j$ is associated with parameters $\theta_j$ and occupation $\tau_j$. Let $\boldsymbol{\tau}$ be the vector of all state occupations and $\boldsymbol{\theta}$ be the vector of all state parameters. The trajectory-state assignments are represented as a binary $N$-by-$K$ matrix $\mathbf{Z}$. The number of states, $K$, is assumed to be known and constant. Our goal is to infer the posterior distribution $p(\mathbf{Z}, \boldsymbol{\tau}, \boldsymbol{\theta} \mid \mathbf{X})$.

The denominator in 1 is intractable for this model, so we choose to construct a tractable approximation $q(\mathbf{Z}, \boldsymbol{\tau}, \boldsymbol{\theta})$ to the posterior. As the criterion for choosing $q$, we seek the $q$ that maximizes the *evidence lower bound* (ELBO):

$$L[q] = -\int q(\mathbf{Z}, \boldsymbol{\tau}, \boldsymbol{\theta}) \log \left[ \frac{q(\mathbf{Z}, \boldsymbol{\tau}, \boldsymbol{\theta})}{p(\mathbf{X}, \mathbf{Z}, \boldsymbol{\tau}, \boldsymbol{\theta})} \right] \mathrm{d}\mathbf{Z} \, \mathrm{d}\boldsymbol{\tau} \, \mathrm{d}\boldsymbol{\theta} \tag{13}$$

where $p(\mathbf{X}, \mathbf{Z}, \boldsymbol{\tau}, \boldsymbol{\theta})$ is the joint probability of all the variables in the mixture model 2.

We can motivate the choice of the ELBO as the "merit function" by appealing to the Kullback-Leibler divergence. Substituting the identity $p(\mathbf{X}, \mathbf{Z}, \boldsymbol{\tau}, \boldsymbol{\theta}) = p(\mathbf{X}|\mathbf{Z}, \boldsymbol{\tau}, \boldsymbol{\theta}) p(\mathbf{Z}, \boldsymbol{\tau}, \boldsymbol{\theta})$ into 13 and rearranging, we have

$$L[q] = \log p(\mathbf{X}) - \text{KL}\left(q(\mathbf{Z}, \boldsymbol{\tau}, \boldsymbol{\theta}) || p(\mathbf{Z}, \boldsymbol{\tau}, \boldsymbol{\theta}|\mathbf{X})\right)$$

where the last term is the Kullback-Leibler (KL) divergence between the true posterior and the approximation. Since $\mathrm{KL}(q||p) \geq 0$ with equality holding only if $q = p$, maximization of $q(\mathbf{Z}, \boldsymbol{\tau}, \boldsymbol{\theta})$ corresponds to minimization of the KL divergence between our approximation and the true posterior. $L[q]$ also provides a lower bound on the log evidence, a property used to select between competing models in approaches such as the vbSPT method [1].

#### C.2 Approximative posterior

How do we specify a functional form for $q$? As it turns out, a tractable form can be induced with a single mean field approximation:

$$q(\mathbf{Z}, \boldsymbol{\tau}, \boldsymbol{\theta}) \approx q(\mathbf{Z}) q(\boldsymbol{\tau}, \boldsymbol{\theta}) \tag{14}$$

The accuracy of the VB approach depends entirely on how good this approximation is.

Under 14, a solution that maximizes $L[q]$ is given by the system of equations

$$\begin{aligned} \log q(\mathbf{Z}) &= \mathbb{E}_{\boldsymbol{\tau}, \boldsymbol{\theta}} \left[ \log p \left( \mathbf{X}, \mathbf{Z}, \boldsymbol{\tau}, \boldsymbol{\theta} \right) \right] + \text{constant} \\ \log q(\boldsymbol{\tau}, \boldsymbol{\theta}) &= \mathbb{E}_{\mathbf{Z}} \left[ \log p \left( \mathbf{X}, \mathbf{Z}, \boldsymbol{\tau}, \boldsymbol{\theta} \right) \right] + \text{constant} \end{aligned} \tag{15}$$

where the constants are chosen so that the factors $q(\mathbf{Z})$ and $q(\boldsymbol{\tau}, \boldsymbol{\theta})$ are normalized. Each expectation is evaluated with respect to the other factor of $q$. VB algorithms often work by iterating between the two expectations, using the previous iteration for each factor of $q$ to evaluate the expectations for the current iteration of the other.

For our trajectory mixture model, we have

$$p \left( \mathbf{X}, \mathbf{Z}, \boldsymbol{\tau}, \boldsymbol{\theta} \right) \propto \prod_{i=1}^{N} \prod_{j=1}^{K} \left( \tau_j f_X(X_i \mid \theta_j)^{Z_{ij}} \right) \prod_{j=1}^{K} \tau_j^{\frac{\alpha}{K} - 1} p(\boldsymbol{\theta})$$

Substituting into the second equation in 15 and using linearity of expectation, we have

$$\begin{aligned} \log q(\boldsymbol{\tau}, \boldsymbol{\theta}) = \sum_{j=1}^{K} \left( \frac{\alpha}{K} - 1 + \sum_{i=1}^{N} \mathbb{E}_{\boldsymbol{Z}} \left[ Z_{ij} \right] \right) \log \tau_j \\ + \sum_{j=1}^{K} \left( \log p(\theta_j) + \sum_{i=1}^{N} \mathbb{E}_{\boldsymbol{Z}} \left[ Z_{ij} \right] \log f_X(X_i | \theta_j) \right) + \text{constant} \end{aligned}$$

The dependences on $\boldsymbol{\tau}$ and each $\theta_j$ occur in separate factors. As a result, $q$ further factors into

$$q(\mathbf{Z}, \boldsymbol{\tau}, \boldsymbol{\theta}) = q(\mathbf{Z}) q(\boldsymbol{\tau}) \prod_{j=1}^{K} q(\theta_j)$$

Analyzing the dependence on $\boldsymbol{\tau}$, we see that the factor $q(\boldsymbol{\tau})$ assumes the form of a Dirichlet distribution:

$$q(\boldsymbol{\tau}) = \text{Dirichlet}\left(\frac{\alpha}{K} + \sum_{i=1}^{N} \mathbb{E}_{\mathbf{Z}}\left[Z_{i,1}\right], ..., \frac{\alpha}{K} + \sum_{i=1}^{N} \mathbb{E}_{\mathbf{Z}}\left[Z_{i,K}\right]\right)$$

Since $\frac{\alpha}{K} + \sum\limits_{i=1}^{N} \mathbb{E}_{\mathbf{Z}}\left[Z_{ij}\right]$ is the expected number of trajectories assigned to state $j$ (with a contribution from the pseudocounts $\alpha/K$), we can see that this posterior "counts" by trajectories. Counting by trajectories leads to systematic state biases in the presence of defocalization because fast states produce many short trajectories while slow states produce a few long trajectories (Appendix B).

To correct this, we can modify $q(\boldsymbol{\tau})$ so that it counts by jumps rather than trajectories, which is more stable to the effects of defocalization. Letting $L_i$ be the number of jumps in trajectory $i$, we write

$$\begin{aligned} q(\boldsymbol{\tau}) &= \text{Dirichlet}\left(n_1, ..., n_K\right) \\ n_j &= \frac{\alpha}{K} + \sum_{i=1}^{N} \frac{dL_i}{2} \mathbb{E}_{\mathbf{Z}}\left[Z_{ij}\right] \end{aligned}$$

(We can obtain this in a less *ad hoc* fashion by modifying the mixture model 2 so that $i$ refers to the $i^{\text{th}}$ jump in the dataset, rather than the $i^{\text{th}}$ trajectory, while continuing to impose the constraint that all jumps in a trajectory originate from the same diffusive state.)

While this is an improvement, it still does not account for the fraction of jumps that land outside the focal volume, which is variable between states with different diffusion coefficients. Appendix B examines this problem in detail; we use these results to derive a full correction later in this section.

With $q(\boldsymbol{\tau})$ in hand, next we turn to $q(\mathbf{Z})$. Evaluating the first factor in 15 and omitting some algebra,

$$\begin{aligned} q(\mathbf{Z}) &= \prod_{j=1}^{K} \prod_{i=1}^{N} r_{ij}^{Z_{ij}} \\ r_{ij} &= \frac{\rho_{ij}}{\sum\limits_{k=1}^{K} \rho_{ik}} \\ \log \rho_{ij} &= \mathbb{E}_{\boldsymbol{\tau}}\left[\log \tau_j\right] + \mathbb{E}_{\boldsymbol{\theta}}\left[\log f_X(X_i|\theta_j)\right] \end{aligned}$$

Since $\boldsymbol{\tau}$ is a Dirichlet random variable under $q(\boldsymbol{\tau})$, we have the well-known result

$$\mathbb{E}_{\boldsymbol{\tau}}\left[\log \tau_j\right] = \psi(n_j) - \psi\left(\sum_{j=1}^{K} n_j\right)$$

where $\psi(x) = \frac{\mathrm{d}}{\mathrm{d}x} \log \Gamma(x)$ is the digamma function. This completes the specification of $q(\mathbf{Z})$.

In order to derive $q(\boldsymbol{\theta})$, first we need to specify the specific form for the trajectory likelihood function $f_X(x|\theta)$. If we use the gamma approximation to the RBME likelihood (equation 11), then each state is parametrized by a single value $\theta_j = 4(D_j \Delta t + \sigma_{\text{loc}}^2)$. Choosing a conjugate inverse gamma prior $\theta_j \sim \text{InvGamma}\left(\alpha/K, \beta_0\right)$, then $q(\theta_j)$ takes on the form of an inverse gamma distribution.

Putting the factors together, the approximative posterior is

$$\begin{aligned} q(\mathbf{Z}) &= \prod_{j=1}^{K} \prod_{i=1}^{N} r_{ij}^{Z_{ij}} \\ q(\boldsymbol{\tau}) &= \text{Dirichlet}\left(n_1, ..., n_K\right) \\ q(\theta_j) &= \text{InvGamma}\left(n_j, \beta_j\right) \end{aligned} \tag{16}$$

where

$$\begin{aligned} r_{ij} &= \frac{\rho_{ij}}{\sum\limits_{k=1}^{K} \rho_{ik}} \\ \log \rho_{ij} &= \mathbb{E}_{\boldsymbol{\tau}}\left[\log \tau_j\right] - S_i \mathbb{E}_{\theta_j}\left[\theta_j^{-1}\right] - \frac{dL_i}{2} \mathbb{E}\left[\log \theta_j\right] \\ n_j &= \frac{\alpha}{K} + \sum_{i=1}^{N} \frac{dL_i}{2} r_{ij} \\ \beta_j &= \beta_0 + \sum_{i=1}^{N} r_{ij} S_i \end{aligned}$$

Here, the relevant expectations are

$$\begin{aligned} \mathbb{E}_{\boldsymbol{\tau}}\left[\log \tau_j\right] &= \psi(n_j) - \psi\left(\sum_{j=1}^{K} n_j\right) \\ \mathbb{E}_{\theta_j}\left[\theta_j^{-1}\right] &= \frac{n_j}{\beta_j} \\ \mathbb{E}_{\theta_j}\left[\log \theta_j\right] &= \log(\beta_j) - \psi(n_j) \end{aligned}$$

This is probably the simplest VB scheme for mixtures of diffusive states in spaSPT data. The algorithm iterates between two steps: (i) evaluate the values of $r_{ij}$ while holding $n_j$ and $\beta_j$ constant, and (ii) evaluate the values of $n_j$ and

$\beta_j$ while holding the values of $r_{ij}$ constant. The posterior mean is

$$\mathbb{E}[\tau_j] = \frac{n_j}{\sum_{k=1}^{K} n_k} \qquad \text{(occupation of state } j\text{)}$$

$$\mathbb{E}[\theta_j] = \frac{\beta_j}{n_j - 1} \qquad \text{(jump variance of state } j\text{)}$$

If the localization error for state $j$ is known, then the diffusion coefficient is

$$\mathbb{E}[D_j] = \frac{1}{\Delta t}\left(\frac{\beta_j}{n_j - 1} - \sigma_{j,\text{loc}}^2\right)$$

where $\Delta t$ is the frame interval.

#### C.3 Correction for defocalization

As mentioned previously, this scheme suffers from a drawback in that it does not account for jumps of trajectories that land outside the focal volume. Suppose that for any state $j$ we have an associated probability $\eta_j$ that a particle inhabiting state $j$ remains within the focal volume until the next frame. (An exact algorithm to compute $\eta_j$ is covered in Appendix B.) If the real occupation of a particular state in the mixture is $p_j$, then the probability to *observe* a jump from this state (which we identify with $\tau_j$) is $\eta_j p_j / \sum_{k=1}^{K} \eta_k p_k$. This can be incorporated in the algorithm above by adding a single final step in which the elements of the posterior Dirichlet distribution $n_1$, ..., $n_K$ are each divided by the respective factor $\eta_j$, so that the final posterior mean occupation for the $j^{\text{th}}$ state is

$$\mathbb{E}[\tau_j] = \frac{n_j/\eta_j}{\sum_{k=1}^{K} n_k/\eta_k}$$

The algorithm represented by eqs. 16 can be considered a Bayesian alternative to least-squares methods for estimating state occupations such as the ones introduced by [13] and [12]. An advantage of this scheme is that it inherits a useful property of Dirichlet priors: when fitting with values of $K$ that are larger than the true number of underlying states, most of the state occupations are driven to zero. In other words, the algorithm favors sparse models with a small number of states that are sufficient to explain the data. This property, a consequence of the "penalty" on model complexity imposed by the prior and incorporated into the ELBO, is quite distinct from the behavior of least-squares or maximum-likelihood approaches that tend to exploit every every parameter of the model, leading to systematic overfitting.

### D Appendix: Subdiffusion with localization error

An attractive possibility for future work is to extend the frameworks of DPMMs and state arrays to non-Brownian motion models. The purpose of this appendix is to highlight the application of state arrays to *subdiffusion*, one of the most commonly reported categories of non-Brownian motion in cells.

Subdiffusion is characterized by mean-squared displacements that scale nonlinearly with time. Specifically, if $F_t$ is the one-dimensional position of a particle at time $t$, then the particle is said to *subdiffuse* if

$$\begin{aligned} \mathbb{E}\left[F_t\right] &= 0 \\ \mathrm{Var}\left(F_{t+\Delta t} - F_t\right) &= 2D\Delta t^{\alpha} \end{aligned} \tag{17}$$

where $\alpha \in (0, 1)$ is an anomaly parameter, and $t \in \mathbb{R}$ (the increment process is stationary). $\alpha = 1$ corresponds to the special case of regular Brownian motion, while $\alpha > 1$ is referred to as *superdiffusion*.

In this section, we begin by deriving the full covariance function for subdiffusion. If the increments are Gaussian distributed, this is equivalent to the tractable *fractional Brownian motion* model. Next, we consider the effect of localization error, much the same way that localization error was added to regular Brownian motion in Appendix A. Finally, we discuss challenges with distinguishing *bona fide* subdiffusion from localization error for short trajectories.

First, we calculate the full covariance function for subdiffusion:

$$\begin{aligned} \mathrm{Cov}\left(F_t, F_s\right) &= \mathbb{E}\left[F_t F_s\right] \\ &= \frac{1}{2}\left(\mathrm{Var}(F_t) + \mathrm{Var}(F_s) - \mathbb{E}\left[\left(F_t - F_s\right)^2\right]\right) \\ &= \frac{1}{2}\left(2Dt^{\alpha} + 2Ds^{\alpha} + 2D\left|t - s\right|^{\alpha}\right) \\ &= D\left(t^{\alpha} + s^{\alpha} - \left|t - s\right|^{\alpha}\right) \end{aligned}$$

(Note that this relies on the stationarity of the motion.)

Since in spaSPT we measure jumps over regular frame intervals of duration $\Delta t$, let $\tilde{F}_n = F_{n\Delta t} - F_{(n-1)\Delta t}$ be the jumps of a subdiffusing particle. We have $\mathbb{E}\left[\tilde{F}_n\right] = 0$ and

$$\begin{aligned} \mathrm{Cov}\left(\tilde{F}_n, \tilde{F}_m\right) &= \mathrm{Cov}\left(F_{n\Delta t}, F_{m\Delta t}\right) - \mathrm{Cov}\left(F_{n\Delta t}, F_{(m-1)\Delta t}\right) \\ &\quad - \mathrm{Cov}\left(F_{(n-1)\Delta t}, F_{m\Delta t}\right) + \mathrm{Cov}\left(F_{(n-1)\Delta t}, F_{(m-1)\Delta t}\right) \\ &= D\Delta t^{\alpha}\left(\left|n - m + 1\right|^{\alpha} + \left|n - m - 1\right|^{\alpha} - 2\left|n - m\right|^{\alpha}\right) \end{aligned}$$

Following Appendix A, we can add positional uncertainty with mean 0 and variance $\sigma^2_{\text{loc}}$ to each point for the final covariance function

$$\begin{aligned}\text{Cov}\left(\tilde{F}_n, \tilde{F}_m\right) = & D\Delta t^\alpha \left(|n-m+1|^\alpha + |n-m-1|^\alpha - 2\,|n-m|^\alpha\right) \\ & + 2\sigma^2_{\text{loc}}\delta(n-m) - \sigma^2_{\text{loc}}\delta(|n-m|-1)\end{aligned} \tag{18}$$

where, as before, $\delta$ is the Dirac delta function. Note that the localization error $\sigma^2_{\text{loc}}$ contributes to both diagonal and off-diagonal terms of the covariance matrix. For $\alpha = 1$, the diffusive terms only contribute to the diagonal. For $\alpha < 1$, *any two distinct jumps in a trajectory have negative covariance.*

If we make the further assumption that the increments are Gaussian-distributed (in which case the covariance function is sufficient to completely determine the stochastic model), then this model is *fractional Brownian motion* [19] with localization error (FBME). In this case, $\alpha = 2H$, where $H$ is the Hurst parameter. The FBME model is attractive in that it the probability of defocalization can be evaluated in a straightforward manner using Monte Carlo integration approaches such as the methods of Alan Genz [18].

We suggest extending the framework of state arrays to FBME by performing inference in the three dimensional parameter space of diffusion coefficient, localization error, and the anomaly parameter $\alpha$. As before, we can marginalize on localization error to obtain a two-dimensional distribution over diffusion coefficient and Hurst parameter. A fundamental challenge for this approach is the close relationship between localization error and subdiffusion in 18. At frame intervals $\Delta t < 10$ ms and with localization error around 40 nm (typical for fast tracking), the magnitude of the contributions of subdiffusion and localization error to the off-diagonal of the covariance matrix are similar. The magnitude of the covariance for terms farther from the diagonal (that is, jumps that do not share a common point in the trajectory), while free of the effects of localization error, becomes vanishingly small except for very small values of $\alpha$. As a result, the problem of disambiguating localization error and subdiffusion is the major challenge to address when applying state arrays to subdiffusion.
